# Telomere maintenance pathway activity analysis enables tissue- and gene-level inferences

**DOI:** 10.1101/2021.02.01.429081

**Authors:** Lilit Nersisyan, Arman Simonyan, Hans Binder, Arsen Arakelyan

## Abstract

Telomere maintenance is one of the mechanisms ensuring indefinite divisions of cancer and stem cells. Good understanding of telomere maintenance mechanisms (TMM) is important for studying cancers and designing therapies. However, molecular factors triggering selective activation of either the telomerase dependent (TEL) or the alternative lengthening of telomeres (ALT) pathway are poorly understood. In addition, more accurate and easy-to-use methodologies are required for TMM phenotyping. In this study, we have performed literature based reconstruction of signaling pathways for the ALT and TEL TMMs. Gene expression data were used for computational assessment of TMM pathway activities and compared with experimental assays for TEL and ALT. Explicit consideration of pathway topology makes bioinformatics analysis more informative compared to computational methods based on simple summary measures of gene expression. Application to healthy human tissues showed high ALT and TEL pathway activities in testis, and identified genes and pathways that may trigger TMM activation. Our approach offers a novel option for systematic investigation of TMM activation patterns across cancers and healthy tissues for dissecting pathway-based molecular markers with diagnostic impact.

## 1 Introduction

Telomeres perform protective functions at the ends of linear eukaryotic chromosomes. They constitute one of the basic molecular factors conditioning the ability of cells to divide. Excessive cell divisions lead to incomplete reconstitution of telomeres resulting in telomere shortening and loss of proper structure of telomeric ends (Blackburn, 1991; Chow et al., 2012; Martĺnez and Blasco, 2015). Highly proliferative cells, such as cancer and stem cells, may utilize mechanisms for preserving the telomere ends despite many rounds of divisions (Greenberg, 2005). These are known as telomere maintenance mechanisms (TMMs, (Hug and Lingner, 2006)). The cells may trigger activation of different TMM pathways, either using the telomerase reverse transcriptase driven synthesis (TEL (Hug and Lingner, 2006; Shay, 2016)), or via DNA break-induced repair (BIR) like processes, also known as alternative lengthening of telomeres (ALT (Cesare and Reddel, 2008; Neumann et al., 2013; Sobinoff and Pickett, 2017; Jia-Min Zhang et al., 2019)). The TEL pathway is more commonly occurring in stem cells and the majority of cancers, while ALT is mostly activated in tumors of mesenchymal and neuroepithelial origin (liposarcomas, osteosarcomas and oligodendroglial gliomas), but can also be found in tumors of epithelial origin (carcinomas of the breast, lung and kidney) (Henson et al., 2002). Some cancers such as neuroblastomas and liposarcomas, do not show evidence for activation of any of the two TMM pathways, exhibiting the ever shorter telomeres phenotype (Costa et al., 2006; Dagg et al., 2017). Finally, some indications have recently prompted that certain cancer entities (liposarcoma and other sarcomas, some tumor types) might also have both of the TMM pathways activated (Costa et al., 2006; Gocha et al., 2013).

Current experimental assays for TMM phenotyping have several shortcomings. For example, the telomeric-repeat amplification protocol (TRAP) assay for estimating telomerase activity is not very sensitive and time and resource consuming (Fajkus, 2006). Assays to measure ALT activity are based on the assessment of chromosomal and/or cellular markers, such as C-circles, ALT-associated nuclear bodies or heterogeneous distributions of telomere length (Pickett and Reddel, 2015; Jia-Min Zhang et al., 2019), which all are usually observed in ALT-type cancers. However, recent studies strongly suggests that none of these markers alone is sufficient to define the ALT status of a cell (Jia-Min Zhang et al., 2019). Attempts to use gene expression signatures for classification of TMM mechanisms have been made (Lafferty-Whyte et al., 2009). However, those signatures are applicable to specific data (Lafferty-Whyte et al., 2009) and they do not provide mechanistic details about TMM pathway activation.

Here we were set to develop a complementary approach to TMM detection that utilizes widely available gene expression data in combination with the molecular interaction topologies in the TMM pathways. Establishment and analysis of TMM pathway topologies is not a trivial issue, because there is no holistic understanding of the functional context of the molecular factors, of their interactions and of the mechanisms triggering TMM activation. Previous research has identified transcriptional regulators of the assembly of the telomerase complex (Yuan et al., 2019), however, how the enzyme components are processed and brought together (Schmidt and Cech, 2015), how the enzyme is recruited to the telomeres and what promotes final synthesis (Chen et al., 2012), is largely not clear and scattered throughout the literature in the best case.

Even less is known about regulation of ALT on a gene level. Although it is considered as a break-induced repair (BIR)-like process, some of the usually accepted BIR factors are not always involved (Jia-Min Zhang et al., 2019). We previously developed a TMM-pathway approach under consideration of TEL and ALT and applied it to colon cancer (Nersisyan et al., 2019). However, overall there is no pathway representation, neither of TEL, nor of ALT TMM, which has been proven in a wider context of cells and/or tissues.

In the first part of the manuscript, we show how gene expression data can be used for TMM phenotyping making use of the TEL and ALT pathways, which were constructed based on available knowledge about molecular factors and interactions of TMM by further developing our previous work (Nersisyan et al., 2019). For demonstration we have analyzed available gene expression data on different cancers with independent experimental TMM annotations (Lafferty-Whyte et al., 2009). In the second part, we apply comprehensive bioinformatics analyses to discover details of TEL and ALT activation in healthy human tissues.

## 2 Results

### 2.1 Literature based reconstruction of telomere maintenance pathways

In order to address the current lack of signalling pathway representations of telomere maintenance mechanisms (TMMs), we have performed a literature search to identify genes involved in TEL and/or ALT TMMs and to define their interaction partners and functional role (Figure 1). Overall, we identified 38 (ALT) and 27 (TEL) genes derived from 19 and 13 references, respectively (Tables 1 and 2). We have considered interactions among these genes in terms of pathway topologies and took into account complex formation and other molecular events with possible impact for TEL or the ALT TMM in order to describe pathway activation in time and space (Figure 1).

**Table 1.**
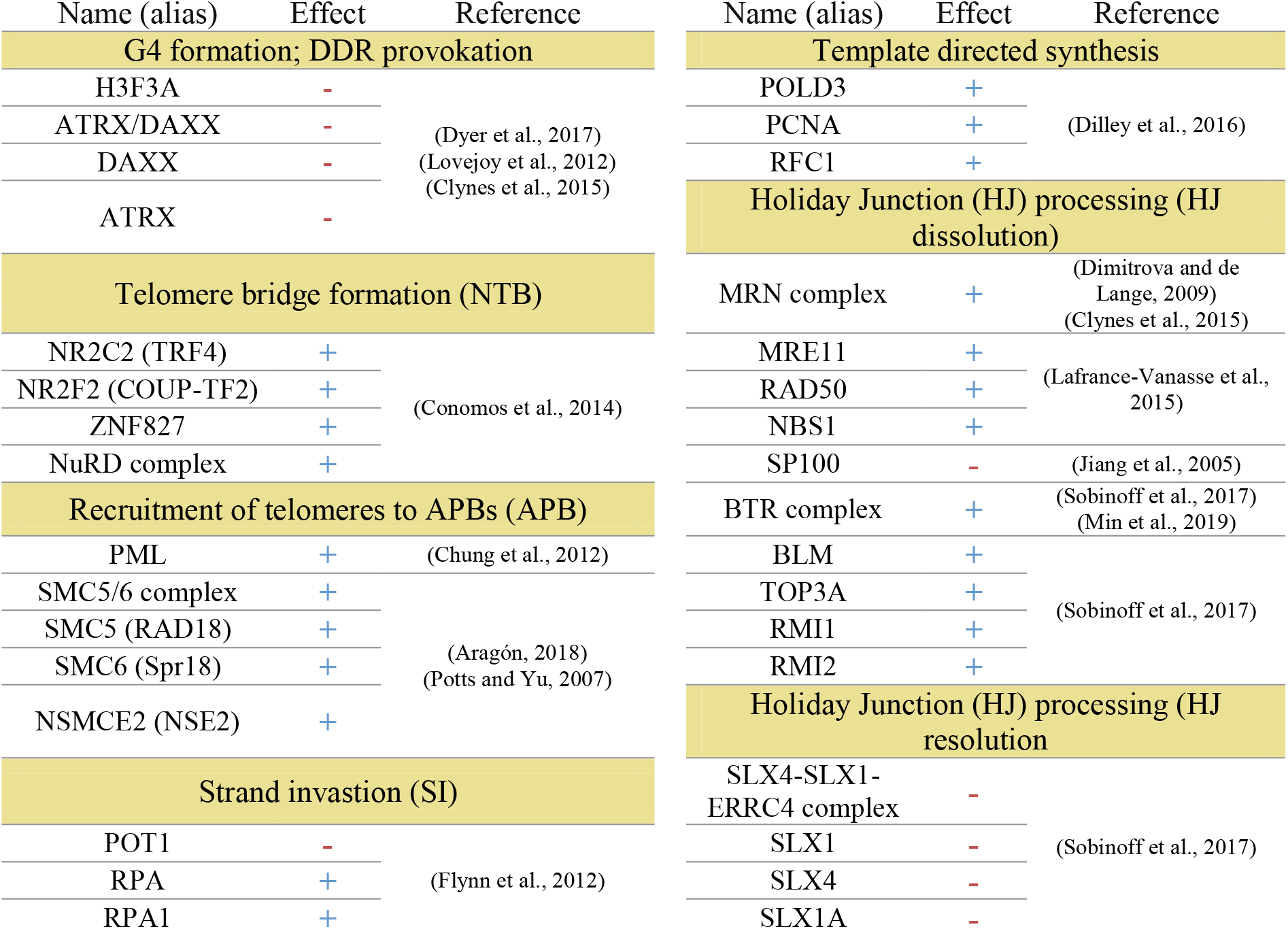
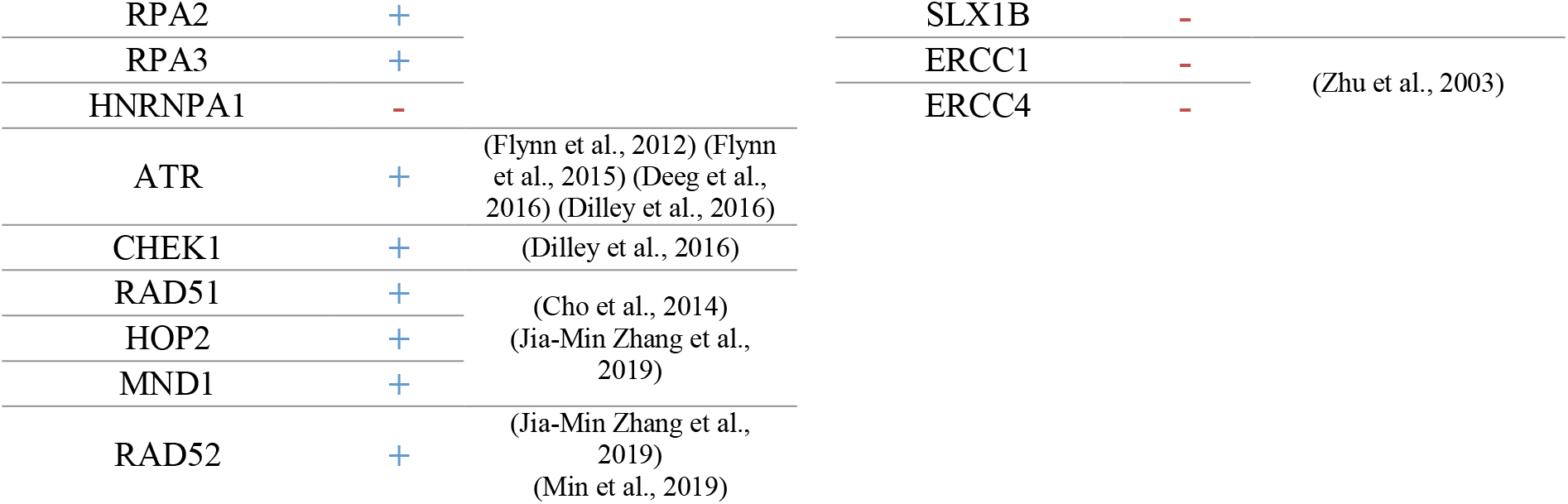
The ALT pathway nodes with activating (+) or inhibiting (-) effects.

**Table 2.**
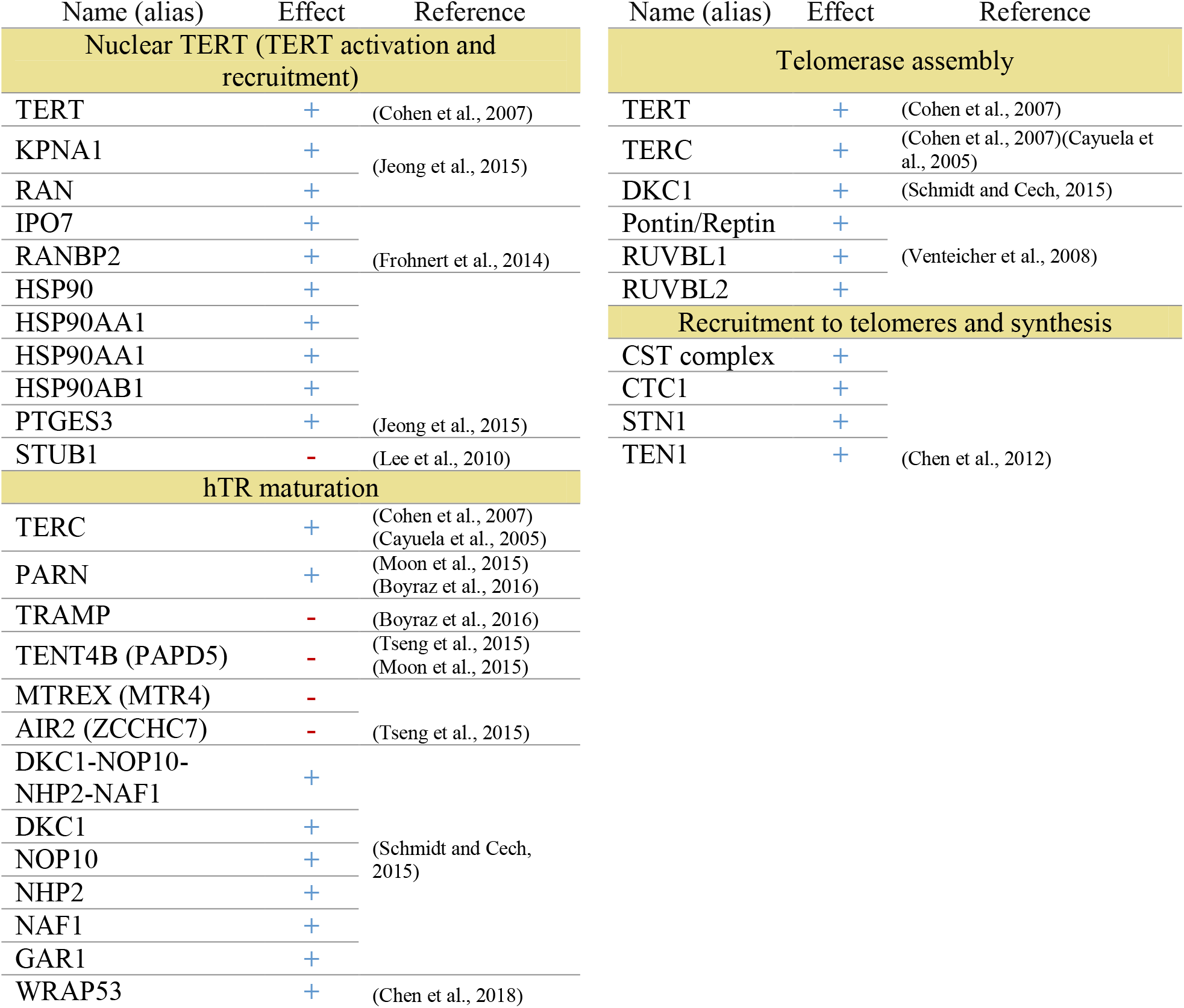
The TEL pathway nodes with activating (+) or inhibiting (-) effects.

**Figure 1.**
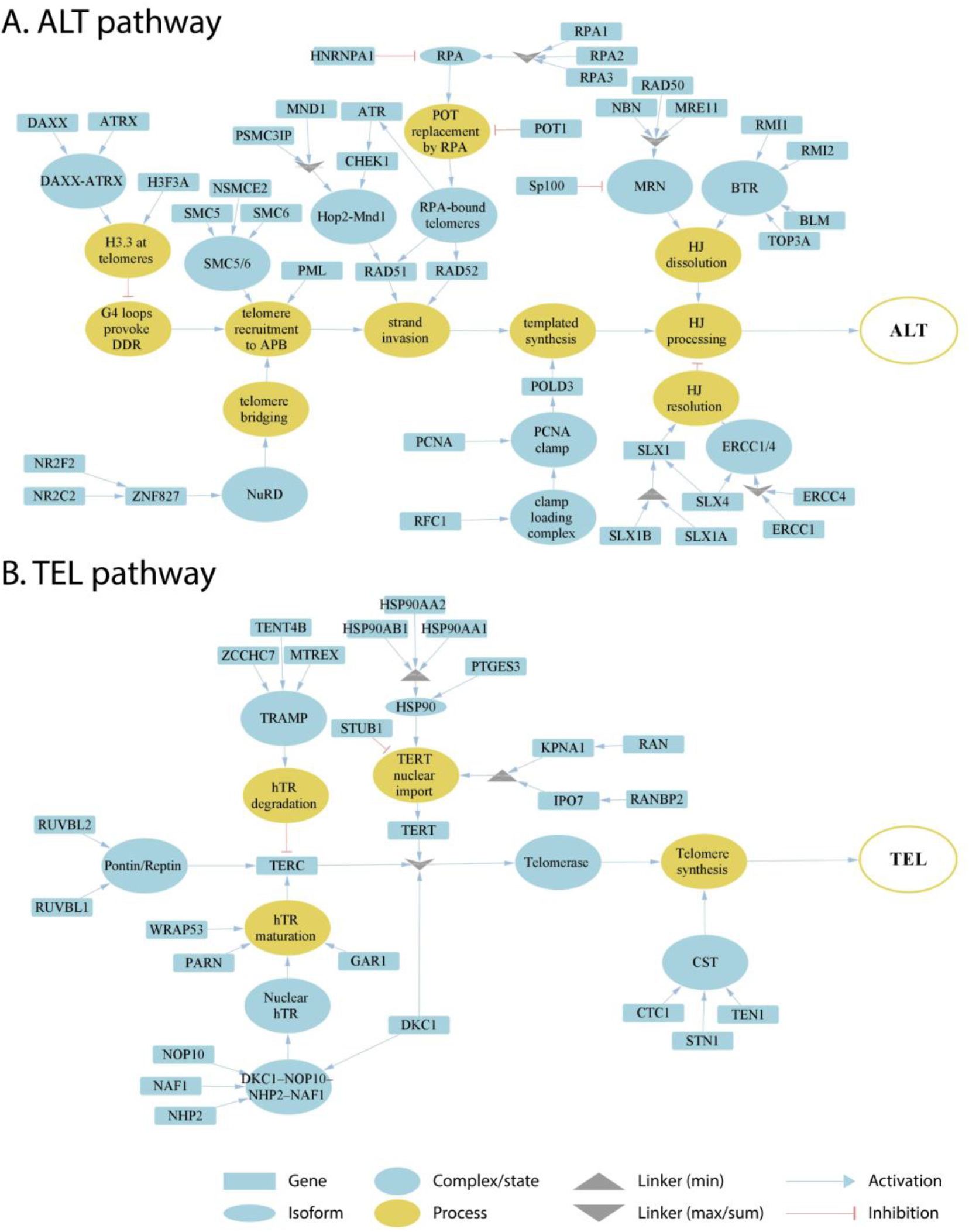
Literature based reconstruction of the ALT and the TEL pathways of telomere maintenance. The ALT and TEL pathways include 37 and 26 genes based on 19 and 13 unique citations, respectively. The events leading to telomere maintenance in each pathway converge at respective final sink nodes (open circles at the right). Types of nodes and edges are defined in the figure. Linker nodes apply different operators for describing signal transduction by taking minimum or maximum values of all the input signals or their sum (see legend in the figure).

Hereby, the ALT pathway is represented through a series of branches describing DNA damage and assembly of APB bodies, which then promote separation of one of the telomere strands and invasion to another telomeric template from sister chromatids, other chromosomes or extra-chromosomal telomeric sequences, followed by DNA polymerase delta assisted telomere synthesis and ultimate processing of Holliday junctions, which is formed during the strand invasion (Figure 1A) (Henson et al., 2002; Pickett and Reddel, 2015; Jia-Min Zhang et al., 2019; Sobinoff and Pickett, 2020). The TEL pathway describes expression, post-transcriptional modifications, recruitment and assembly of different components of the telomerase complex, namely hTERT, hTR and dyskerin, followed by the formation of a catalytically active telomerase complex, its recruitment to telomeres and telomere synthesis by telomerase and DNA polymerase alpha (Figure 1B)(Hug and Lingner, 2006; Tseng et al., 2015; Rice and Skordalakes, 2016).

### 2.2 TMM pathway activities are supported by experimental test assays

For assessment of TEL and ALT TMM pathway activities in a given sample we use gene expression data and the pathway signal flow (PSF) algorithm as implemented in Cytoscape (apps *PSFC* and *TMM*; see methods section for details). The algorithm computes the PSF score in each of the pathway nodes by considering signal propagation through all upstream activating and inhibiting interactions and complex and linker node types making use of the fold change (FC) expression values of the involved pathway genes with respect to their mean expression in the respective data set. The PSF score thus reflects the activity of all upstream events and of their topology in contrast to gene set overexpression measures often used alone for functional assessment (Hakobyan et al., 2016; Nersisyan et al., 2017). The PSF scores of the final sink nodes then estimate the overall activity of the TEL and the ALT pathway, respectively.

Application to two publicly available microarray gene expression datasets from cell lines and liposarcoma tissues delivers an ALT/TEL-PSF data couple for each sample, which is then plotted into an ALT-versus-TEL PSF coordinate systems (Figures 2A, B). Each sample is color-coded according to its assignment to TEL^+^/TEL^−^ or ALT^+^/ALT^−^ or double negative ALT^−^/TEL^−^ phenotypes which were determined by independent experimental assays alongside with the gene expression measurements. Using support vector machine learning, TEL positive and negative samples were separated by a vertical line, and ALT positive and negative samples by a horizontal line in both data sets (Figure 2A, B).

**Figure 2.**
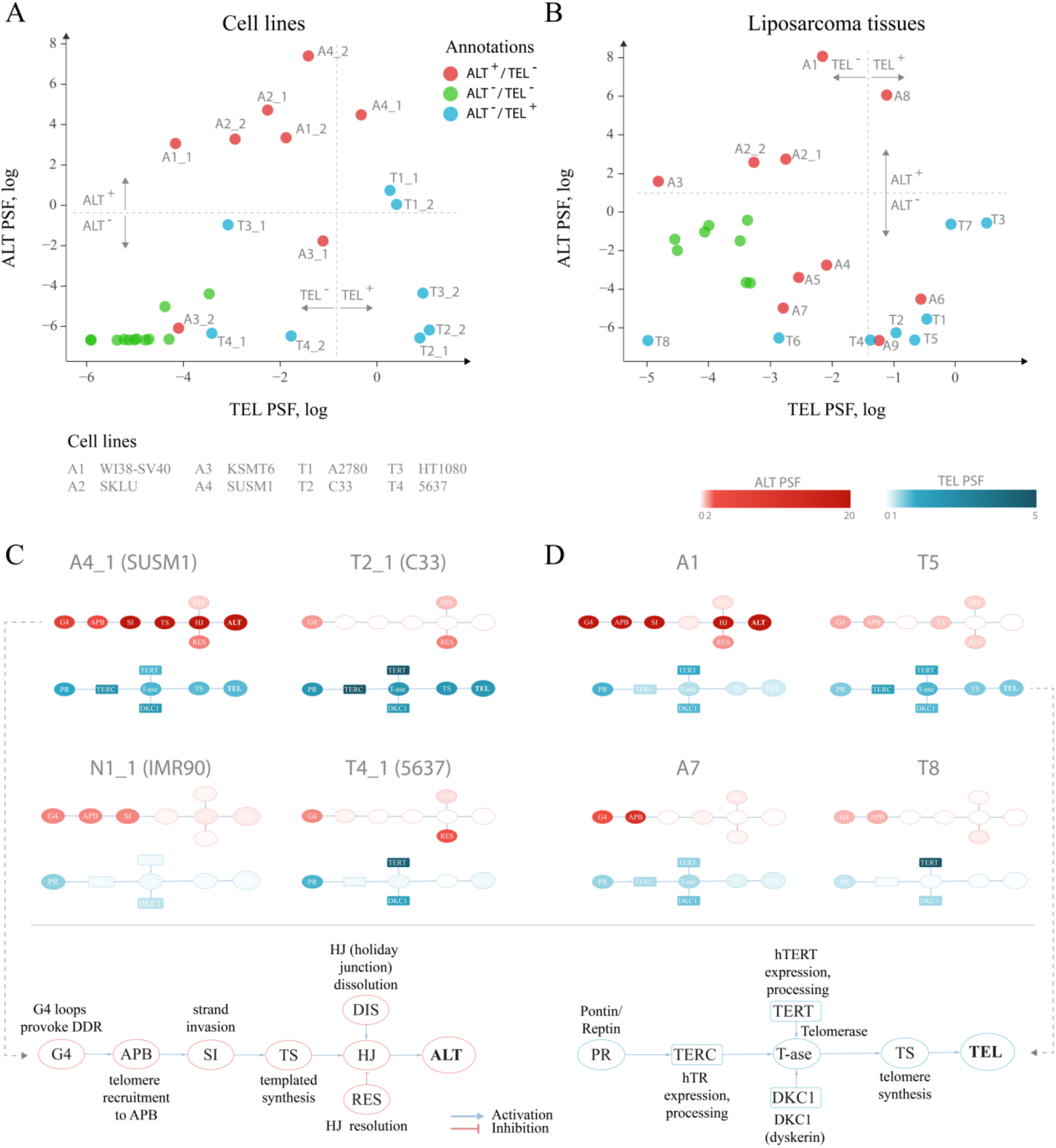
TEL and ALT pathway activity plots for cell lines and liposarcoma tissues. **A**. Cell lines and hMSCs plotted according to the TEL PSF (x axis) and ALT PSF (y axis) values. Color coding corresponds to experimental annotation of TMM states. Horizontal and vertical dash lines separate ALT^+^ from ALT^−^ and TEL^+^ from TEL^−^ experimentally annotated samples based on support vector machine classification on the ALT and the TEL PSF values. Technical replicates are distinguished with _1 and _2 suffices. **B**. Similar representation for liposarcoma tissues and hMSCs. **C-D**. Examples of pathway activation patterns of some cell lines (C) and liposarcoma tissues (D). PSF values of process nodes are indicated on a light-to-dark color scale. Node abbreviations are explained in the bottom schemes. Pathway activity patterns for all the samples are shown in Supplementary Figures S2 and S3.

Detailed inspection revealed that all double negative samples (green circles) indeed locate in the ALT^−^ /TEL^−^ quadrant formed by the perpendicular lines of our classification scheme thus indicating perfect agreement with experimental assays. For single positive ALT^+^/TEL^−^ (red) and ALT^−^/TEL^+^ (blue) samples one finds their accumulation in the respective top-left and down-right quadrants, respectively, as expected. A certain fraction of theses phenotypes is found ‘displaced’ in the left-down and top-right quadrants assigning them to double negative and double positive TMM cases, respectively. For some samples, we observed discordance in TMM PSF values between technical replicates, which was also noticeable on the level of gene expression (Supplementary data 2), suggesting possible technical issues during microarray processing. Overall, we obtained 80-85% agreement between our computational assessment of TMM activity as ALT or TEL single positive and full agreement with double negative samples and the experimental annotations taken from the original publication.

### 2.3 Analysis of pathway activation patterns at single gene and single-sample resolution

For a better understanding of the particular reasons of the diversity observed in the ALT/TEL plots we visualized pathway activity patterns for selected samples in Figure 2C, D by colouring the major event nodes in light-to-dark blue (TEL PSF) or red (ALT PSF) (the nodes were annotated in the part Figure 2C, D; the full gallery of pathway activation patterns for all samples is provided in Supplementary Figures S2 and S3).

SUSM1 (A4), a transformed ALT-activated (according to experimental assignment) cell line derived from fetal liver, shows activation of all the nodes involved in TEL and ALT (Figure 2C), which pushes one replicate to the upper right quadrant corresponding to the double positive TEL^+^/ALT^+^ pathway phenotype. The C33 cell line (T2), derived from cervical squamous cell carcinoma, shows clear activation of the major TEL pathway nodes, while the main ALT events, such as telomere recruitment to APB, strand invasion and telomere synthesis are suppressed, pushing it to the TEL^+^/ALT^−^ quadrant in agreement with experimental assignment (Figure 2C). The bladder carcinoma cell line 5637 (T4), experimentally assessed to have high telomerase activity, showed relatively low TEL PSF values. As seen in Figure 2C, the expression and processing of the two telomerase complex subunits hTERT (TEL pathway branch TERT) and dyskerin (branch DKC1) were highly activated in this cell line, however the expression of the factors processing the RNA template hTR was low, thus being a bottleneck for the telomerase complex formation, explaining the observed discrepancy. We could also identify the genes responsible for this, as described in detail below.

Looking at the patterns of the A7 sample in the dataset of liposarcoma tissues (Figure 2D), we observed high activity of the APB branch of the ALT pathway, indicating that both the computational annotation and the experimental assay point on accumulation of APB bodies in this tissue. However, as the expression of downstream factors in the ALT pathway was low (Figure 2D), it compromised the ultimate activation of this pathway. This result supports recent studies showing that the mere presence of APB bodies doesn’t necessarily lead to the activation of ALT (Jia-Min Zhang et al., 2019). Similar branch-activation patterns were observed for the A6 and A9 tissue samples (Supplementary Figure S3). It is important to note that these samples had high TEL pathway activity, which is in line with previous observations of high false-positive rate of the APB assay associated with overexpression of *TERC* or *TERT* (Henson and Reddel, 2010). In summary, inspection of the PSF patterns along the pathways identifies genes and branches contributing to activation or deactivation of ALT and TEL with single sample resolution. More information regarding gene-level activation patterns can be explored in the full pathway PSF activation patterns (Supplementary data 3).

### 2.4 Partial influence (PI) analysis identifies gene-specific triggers of pathway activities

As a second additional option of extracting gene level information responsible for pathway activation changes we analyzed the partial influence (PI) of each gene (Material and methods section). PI enables understanding of which genes act as triggers to activate or to deactivate selected nodes in the pathways. Genes, increasing or decreasing the PSF of the target node have positive or negative PI’s respectively. Activating nodes with log fold change (FC) expression above or below zero in a given sample will thus have a positive or negative PI respectively. Nodes with inhibitory effect, on the other hand, will have the reverse association with PI (Figure 3A).

**Figure 3.**
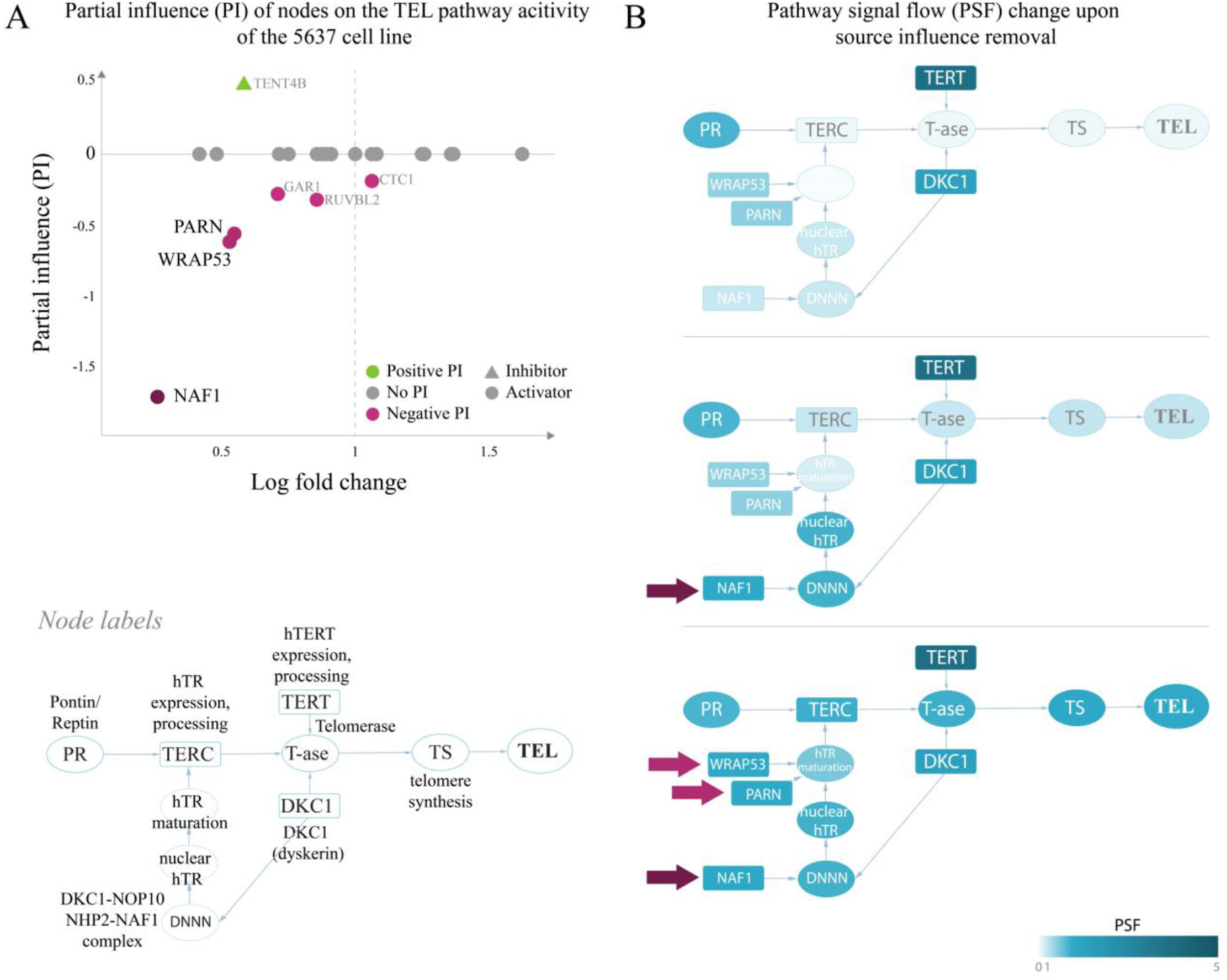
Partial influence of nodes on the TEL pathway activity of the 5637 cell line. **A**. Partial influence (PI) of each node is computed as the difference of the TEL node PSF value when the node is set to a fold change (FC) value of 1. In the first replicate of the 5637 cell line, the genes involved in maturation of the telomerase RNA component hTR (*NAF1, WRAP53* and *PARN*), have the largest negative influence on the TEL pathway activity. Low expression of these genes leads to inactivation of the TERC branch, which is the bottleneck in telomerase complex formation (**B**). The pathway becomes activated when the influence of *NAF1* (**C**) or *NAF1, WRAP53 and PARN* (**D**) is removed by setting their FC to 1 (see Methods).

We have examined the PIs in the TEL pathway of the 5637 (T4) bladder carcinoma cell line (Figure 2A), where branch-level analyses showed suppression of the hTR maturation branch (Figure 2C). The results of the PI analysis show that the genes encoding the hTR maturation factors *NAF1, WRAP53* and *PARN* (Figure 3) were responsible for low activity of the hTR branch in this sample.

Indeed, bringing the relative expression of *NAF1* to a fold change value of 1 (Figure 3B) was sufficient to push the activity of the TEL pathway over the threshold for classifying it as TEL^+^ (log PSF of 0). It is important to mention that the used microarray gene expression datasets did not contain expression values for the *TERC* gene itself, and only the expression of hTR processing factors contributed to the hTR branch in this case.

A similar branch activation and PI pattern was observed for the liposarcoma tissue T8, where the TMM-PSF diverged from the experimental assay results (Figure 2B, Supplementary Figure S6). In the other misclassified sample (T6), we observed high PSF activity of the telomerase complex, however, the low TEL pathway activity was driven by low expression of STN1, which stimulates polymerase alpha in synthesis of the complementary telomeric strand after the action of the telomerase complex (Supplementary Figure S6).

PI analysis also identified that various genes were responsible for activation of the APB branch of the ALT pathway in the A6 and A7 liposarcoma tissues, while down-regulation of RAD51 and CHEK1 led to suppression of the downstream strand invasion events leading to low ALT PSF activity in both samples (Supplementary Figure S5). Hence, PI analysis extracts genes which act as triggers for switching TMM on or off with possible impact for altering between ALT and TEL and *vice versa*.

### 2.5 Comparison with TelNet genes

Our curated TMM-pathway approach considered in total 63 genes extracted from recent publications (see above and Materials and Methods section). As an alternative option we made use of TelNet data base (Braun et al., 2018) which collected 2,094 genes with impact for telomere biology and extracted 336 genes annotated as ALT- or TEL-associated (Supplementary Table S3). Separate hierarchical clustering of the expression values of the TEL- and ALT-genes in the cell line and liposarcoma samples analyzed above, well separates double negative ALT^−^/TEL^−^ from the single positive ALT and TEL samples on one hand, and also the latters each from another. Between 82% and 85% of the samples were properly annotated compared with the experimental annotations. Agreement with PSF-based annotations is high (96 and 100%, respectively). Hence, gene clustering and PSF based classifications and experimental annotations are well-aligned (Figure 4), and those samples that were misclassified by the PSF algorithm were also misclassified with the TelNet gene set clustering. This simple comparison served as an independent validation for the selection of genes in our TMM-PSF approach using TelNet. Note that the overlap between both collections is 42 genes, meaning that 67% of the TMM-pathway genes are considered in TelNet what we attribute to our more recent curation. Moreover, the TMM pathway approach clearly makes use of a markedly reduced number of genes after strict curation and, moreover, enables topology-based analysis not only to extract genes, but also relevant branches and triggers for switching between TMM pathways.

**Figure 4.**
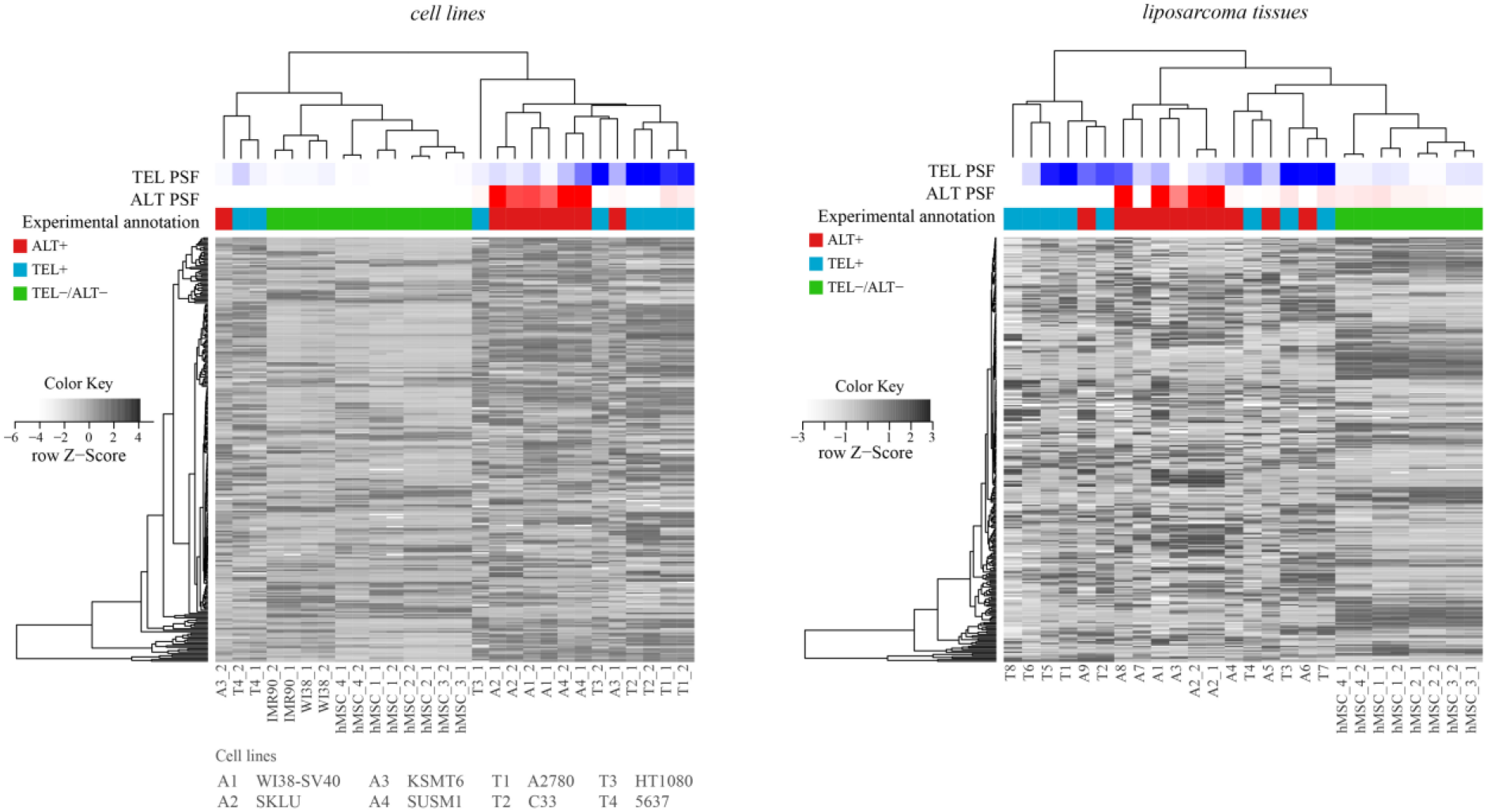
Hierarchical clustering of cell lines and tissues based on expression of genes retrieved from the TelNet database. TEL and ALT PSF values are depicted with blue and red color gradients, and experimental annotations are drawn below. PSF and TelNet assignments closely agree each with another.

### 2.6 Telomere maintenance in healthy human tissues

We have previously applied our PSF approach to study telomere maintenance states in Lynch syndrome and sporadic colorectal cancer subtypes (Nersisyan et al., 2019). Here we expand this method to evaluate the state of telomere maintenance mechanisms in healthy human tissues.

We estimated TEL and ALT activities in units of PSF across a series of fifteen tissues (overall N = 17-80 samples per tissue taken from donors evenly distributed across age and sex groups) making use of RNA-seq data taken from the GTEx portal (Figure 5). The TEL pathway activity was generally low across the tissues (Figure 5A), while ALT shows slightly enhanced PSF-values (Figure 5B).

**Figure 5.**
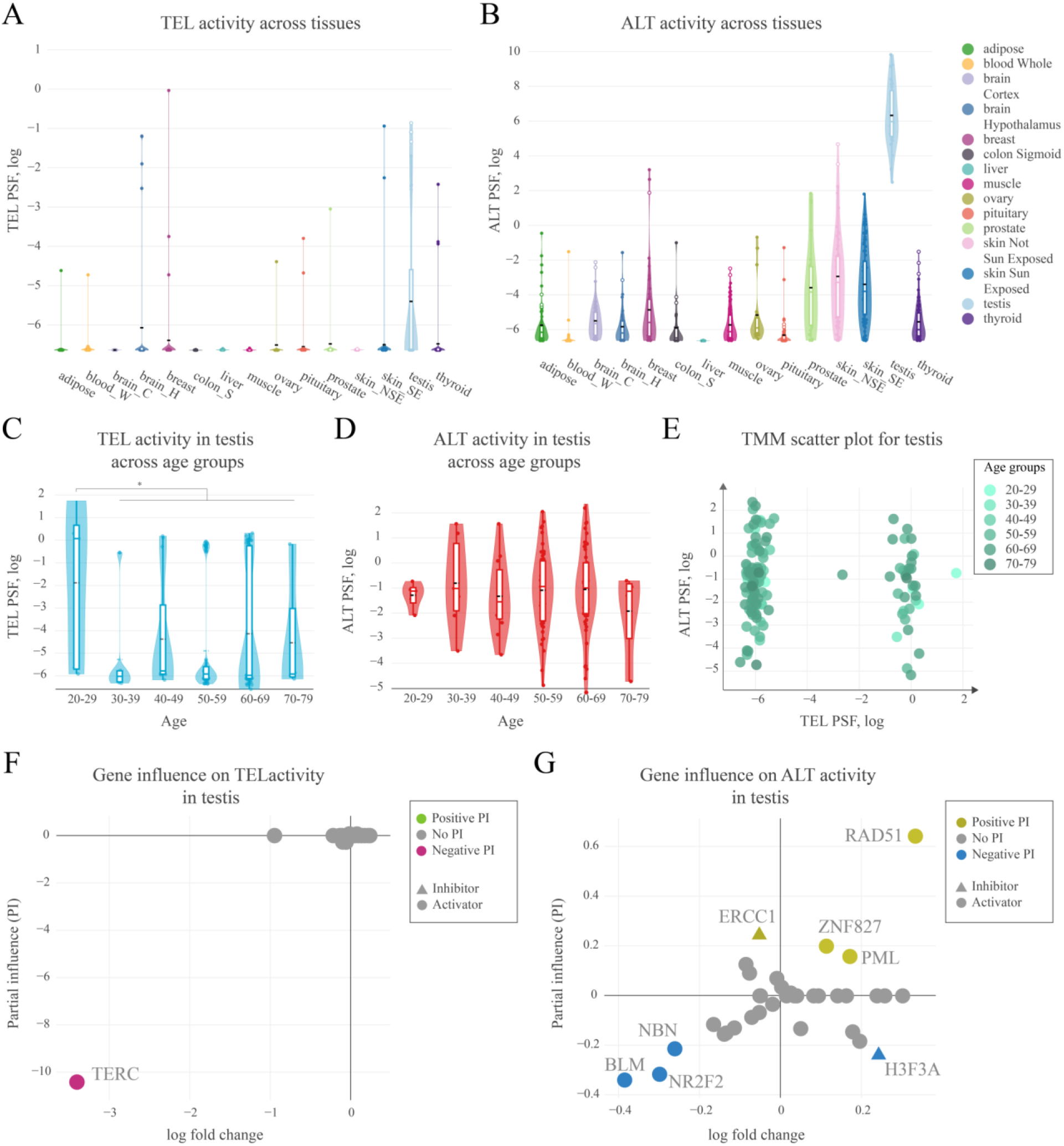
Activity of telomere maintenance mechanisms in healthy human tissues. **A**. TEL pathway activity in healthy human tissues. The black dashed lines represent means, the colored lines represent medians. **B**. ALT pathway activity in human tissues. **C**. TEL pathway activity in testis across age groups. **D**. ALT pathway activity in testis across age groups. **E**. ALT and TEL pathway activity scatter-plot in testis colored by age-groups. **F**. Partial influence (PI) of TEL pathway genes on the pathway activity. **G**. PI of ALT pathway genes on the pathway activity.

Interestingly, testis showed considerably increased activity of ALT consistently in all of the samples, and also activation of TEL, however, only in half of them, while the other half showed virtually deactivated TEL (Figure 5 A, B).

For a more detailed view on the TMM activation patterns in testis, we stratified the PSF-values of the full sample set (N=129) according to the age of the donors (Figure 5 C, D). No clear ageing-trend of TEL and ALT PSF was observed except a shift of the TEL-PSF distribution towards larger values for younger men of age 20-29 years (Mann-Whitney U test *p*=0.01). On the other hand, we observed a clear bi-modal distribution of samples with high or low TEL pathway activities regardless of age (Figure 5 E). The detailed analysis of TEL pathway activation patterns in terms of PI showed that the observed variability was mainly driven by the expression of *TERC* (Figure 5F). This gene was not expressed in half of the testis samples leading to very low TEL activities. Importantly, *TERC* was generally found the main limiting factor for TEL pathway activity across all the tissues studied meaning that low *TERC* expression strongly downscales TEL-PSF.

PI analysis of the ALT pathway in testis mainly implicated the importance of *RAD51*, suggesting RAD51-dependent ALT activation. In the RAD51-dependent pathway, telomere-bound POT1 is replaced by RPA, followed by RAD51 recruitment and strand invasion. Low expression of BLM helicase and of the nuclear receptor NR2F2 potentially limits hyperactivity of ALT in this tissue (Conomos et al., 2014; Sobinoff et al., 2017; Zhang et al., 2021), although it has recently be shown that in some cases NR2F2 may not be directly involved in ALT (Alhendi and Royle, 2020). Interestingly, a recent study of Episkopou *et al*. have identified that the expression of the testis-specific Y-encoded-like protein 5 (*TSPYL5*), a previously unrecognized APB body component, is crucial for survival of ALT^+^ cells, as it protects replaced POT1 from proteasomal degradation (Episkopou et al., 2019). We also found high expression of *TSPYL5* in our testis samples showing slight correlation with ALT activity (Pearson R = 0.3, Supplementary Figure S7). This supports possible involvement of *TSPYL5* in maintaining viability of ALT^+^ cells in healthy human testis.

In summary, our analyses show that both TEL and ALT pathways of telomere maintenance may be activated in human testis. In this tissue, the main driver for ALT pathway activation is *RAD51*, while the limiting factor for TEL pathway activity is the expression of *TERC*, which was observed only in a subset of samples, notably more pronounced in younger subjects.

## 3 Discussion

The activation of TMM mechanism can serve as a phenotypic biomarker for prognostic purposes and for choosing chemotherapies, e.g. by direct targeting of the TMM pathways (Villa et al., 2008; Sugarman et al., 2019; Chen et al., 2020). However, little is known about the molecular triggers leading to activation of telomere maintenance mechanisms via the TEL or the ALT pathways. Some cancer tissues are prone to activation of the ALT pathway, such as the mesenchyme-originating liposarcomas, osteosarcomas and glioblastomas, while others are more permissive of TEL activation (Henson et al., 2002). At the same time, some tumors do not activate neither TEL nor ALT TMM pathways, while others may activate both or switch activation from TEL to ALT or vice versa (Costa et al., 2006; Gocha et al., 2013). The molecular mechanisms behind such cellular decisions are mostly unknown. Owing the role of TMM in cancer prognosis and the promise of telomere-targeting therapies (Villa et al., 2008; Sugarman et al., 2019; Chen et al., 2020) it is of paramount importance to investigate these activation patterns, as well as to come up with better approaches to assess or predict TMM states of tissues and individual cells.

Our study aimed at combining information about molecular factors involved in the TMMs to study the mechanisms of activation of either the TEL or the ALT pathways in cancers and healthy human tissues, and to provide a complementary bioinformatics method for identification of TMM phenotypes from gene expression data. To reach this goal, we have reconstructed TEL and ALT pathways of telomere maintenance taking into account, first of all, comprehensive review and recent original articles of the last three years (Supplementary Tables S1, S2). To the best of our knowledge this is the first attempt of providing a holistic view and quantitative analysis of signalling events involved in TMM.

The TMM pathway topologies proved by quantitative assessment of the TEL and ALT TMMs activity in cancer cell lines and tissues based on two gene expression data taken from a previous study (Lafferty-Whyte et al., 2009). Our pathway approach not only considered the expression values of the genes in each pathway, but also their mutual (activating or inhibiting) interactions, complex formation as well as linking operator nodes all together potentially influencing the final pathway activity states. According to the combinations of the activity of TEL and ALT we have classified the samples into four TMM phenotypes (TEL^−^/ALT^−^, TEL^+^/ALT^−^, TEL^−^/ALT^+^, TEL^+^/ALT^+^) in 80-85% agreement with independent experimental annotations of TMM in the samples. The absence of *TERC* expression data due to the lack of microarray probes for this gene may have limited the accuracy of the TEL pathway activity estimation in part of the samples.

Our TMM-PSF method stands out with a couple of advantages: (a) it helps to easily annotate samples based on ALT/TEL pathway activity values, and (b) it provides molecular details for dissecting the role of genes and sub-events in the overall activation of the pathway. For example, we could show that for some samples despite their low ALT activity, the APB-pathway branch was highly active, which may explain why these samples were detected as ALT positive in the independent APB assay. This is in agreement with recent studies showing that the existence of APBs does not ensure telomere synthesis and many APBs in the cell may lack ALT activity (Jia-Min Zhang et al., 2019). In consequence APB-based ALT tests may lead to false positives in samples with telomerase overexpression (Henson and Reddel, 2010).

It is important to note that there is no gold-standard method for TMM phenotyping of cells/tissues. All currently available experimental assays have their drawbacks (Pickett and Reddel, 2015; Jia-Min Zhang et al., 2019). The activity of the telomerase enzyme is usually assessed by the TRAP assay that measures the amount of DNA synthesized from a telomere-like template *in vitro*. However, it has low sensitivity, is not well suited for single-cell analysis and does not account for downstream processes, such as recruitment of the enzyme to the telomeres and synthesis of the complementary strand (Fajkus, 2006). ALT activity is assessed by assays that measure the abundance of extra-telomeric C-circles or of APB’s, or heterogeneity of telomere length (Sobinoff and Pickett, 2017). However, while these biomarkers may be common in many ALT positive cells/tissues, it has been shown that they alone are neither sufficient nor required for promoting ALT (Jia-Min Zhang et al., 2019). New methods for direct monitoring of telomere elongation in ALT cells are still being adapted (Verma et al., 2018).

There is an urging need for novel assays to detect TMM states that could help in understanding cellular response to chemotherapies and for development of TMM targeted therapies (Villa et al., 2008; Sugarman et al., 2019; Chen et al., 2020; Recagni et al., 2020). Particularly useful will be assays that allow for TMM assessment in single cells. As it has been shown previously, and confirmed in this study, both TEL and ALT pathway may be co-activated in the same tumor. It will be extremely important to assess this issue by single cell transcriptomics and our pathway approach whether those pathways co-exist in the same cell or show mosaic activation in the tissue (Costa et al., 2006; Gocha et al., 2013). Additionally, switching from TEL to ALT as a cellular response to a therapy is also possible (Gan et al., 2002; Bechter et al., 2004; Shay et al., 2012; Recagni et al., 2020). It is possible that the poorer agreement with experimental assays in the liposarcoma tissue samples, compared with the cell lines, was caused by the presence of different cell types in those tissues. In this sense, using RNA-seq gene expression data to assess TMM activity from single cells is a promising future direction.

We have previously applied our approach to dissect molecular factors involved in TMM activation in Lynch syndrome and sporadic colorectal cancer subtypes in order to study association of ALT with microsatellite and chromosomal instability in those cancers (Nersisyan et al., 2019). Here we expanded beyond these studies to investigate TMM activation patterns in healthy human tissues. The experimental assays have so far been used to detect ALT activity in cancers only. Previous studies using telomeric DNA tagging have found evidence for telomere elongation via the ALT pathway in the absence of high telomerase activity in mammalian somatic tissues during early development (Liu et al., 2007; Neumann et al., 2013) The recent study by Novakovic *et al*. has identified elevated TERRA levels, elongated telomeres and some evidence for ALT-specific C-circles in human placenta (Novakovic et al., 2016). The authors note, however, that the amount of C-circles was low compared to ALT positive cancers, suggesting that a mild ALT phenotype may exist in specific placental cells.

Overall, these studies show that more sensitive methods to detect low ALT levels are needed to further investigate TMM states in healthy human tissues with higher resolution. In this study, gene expression datasets from healthy human tissues from the GTEx portal were used to assess TMM activity states. We found high ALT and TEL pathway activities, first of all, in testis. Interestingly, TEL was activated only in a subset of testicular tissues paralleled with marked *TERC* expression, while depleted *TERC* levels lead to low TEL phenotypes. Indications for analogous binary mosaicism effects were reported in previous studies. In humans, *TERC* is mainly expressed in the primary spermatocytes, however at a lower level than in other spermatogenic cells (Paradis et al., 1999). In addition, *TERT* expression also shows mosaic-like regulation in testis, depending on the cells of the tissue or on the stage of spermatogenesis (Ozturk, 2015). Another study has shown that telomere length increases during the development of male germ cells from spermatogonia to spermatozoa, inversely correlated with enzymatic activity of telomerase (Achi et al., 2000). This could provide the link between the observed TEL mosaicism, and activation of the ALT pathway in testis.

The high ALT activity observed in testis was largely conditioned by upregulation of *RAD51*, suggesting that the ALT pathway is activated in a RAD51-dependent, rather than independent manner in testis (Jia-Min Zhang et al., 2019). Interestingly, the expression of the testis-specific Y-encoded-like protein 5 (*TSPYL5*), a recently identified APB body component that is crucial for survival of ALT^+^ cells (Episkopou et al., 2019) was associated with ALT activity in healthy human testis in our study. Finally, our results indicate elevated TMM activity in testis in agreement with the recent study, also performed on GTEx datasets. Accordingly, telomeres are the longest in healthy human testis compared with other tissues and are in weak negative correlation with age (Demanelis et al., 2020).

In summary, we have reconstructed the TEL and ALT TMM pathways from previous literature knowledge. It has been carefully curated relying on reported molecular ingredients contributing to TMM and their interactions. Pathway signal flow activity estimates obtained from gene expression data have been shown to reliably estimate the TMM phenotype as a novel complementary approach to experimental assays. The main advantage of our approach is its “white box” (in contrast to “black box”) nature meaning that the resulting TMM phenotype can be “dissected” at gene level. In other words, we can explore gene-specific effects on sub-processes or events triggering activation of TMM. The method estimates TEL and ALT pathway activity in the same sample, thus enabling to establish its state in a TEL/ALT continuum with impact for investigating co-activation and switching events between the two pathways, e.g., upon cancer treatment and development. Of importance, owing to its sensitivity, it detects subtle activation of TEL and ALT in healthy human tissues. Despite the actuality of our pathway topologies it is important to note that new studies about additional factors affecting TMMs are permanently appearing with possible consequences for the pathways reconstructed here. Notably, the topologies were reconstructed in a generic manner: as the pathways can be activated via different mechanisms, not all the genes are required for the pathway activation in different situation. Future applications will show what branches and components are important in different situations. Single cell transcriptomics is one important field to essentially improve resolution of our TMM pathway method. Expression based methods thus potentially provide an independent and complementary approach to assess the TMM state. Because of the complex nature of TEL and ALT TMM, single gene expression markers or gene set signatures are potentially insufficient in most cases. Our TMM pathway method paves a new way for omics-based evaluation of TMM with a series of applications ranging from cell experiments to cancer diagnostics.

## 4 Materials and methods

### 4.1 Literature search and pathway construction

TEL and ALT pathways presented in this publication were carefully constructed based on comprehensive literature search resulting in 31 publications which provided relevant knowledge about TMM. To find and select publications describing factors involved in the telomere maintenance mechanisms, we searched the PubMed database with terms “telomerase” or “alternative lengthening of telomeres”. The first phase was based on “review” articles published before 2020. These reviews provided the order of molecular processes involved in the pathways and thus their basic topology. The TMM genes mentioned in the review articles were chosen and then used in an extended search of the form [gene or protein] AND (“telomerase” OR “alternative lengthening of telomeres”). We also included genes not yet mentioned in review articles, by repeating the initial search with “telomerase” or “alternative lengthening of telomeres”, concentrating on the original research articles of the last three years (2017-2020).

The articles were read in chronological order and in case of current consensus about the functional role of the mentioned genes they were included in the respective pathway, otherwise they were ignored. To ensure quality, we confirmed each interaction by inter-researcher agreement between the two authors of this manuscript that have curated the pathways. The present version of ALT and TEL pathways (TMMv2.0) is based on a previous version (TMMv1.0) (Nersisyan, 2017; Nersisyan et al., 2019), and makes use of an improved topology, branch and interaction structure according to updated literature knowledge. The network in .xgmml format may be accessed from Supplementary data 4.

### 4.2 Data sources and preprocessing

For approval of ALT and TEL TMM pathways we made use of gene expression data on cell lines and liposarcoma tissues taken from the study by Lafferty-Whyte *et al* (Lafferty-Whyte et al., 2009) which were annotated as double negative ALT^−^ /TEL^−^ (ALT and TEL inactive) or single positive ALT^+^/TEL^−^(ALT active) or ALT^−^/TEL^+^ (TEL active) using independent experiments. The gene expression matrix files were downloaded from the Gene Expression Omnibus (GEO) repository (accession GSE14533). This dataset contains microarray gene expression profiling data for ten cell lines cultured from different tissues, and seventeen liposarcoma tumor samples along with four human Mesenchymal Stem Cells (hMSC) samples isolated from the bone marrow of healthy individuals. ALT activation was assessed by the presence of ALT-associated promyelocytic leukemia bodies (APBs) (Costa et al., 2006; Cairney et al., 2008b), while TEL activation was measured using the telomeric-repeat amplification protocol for telomerase activity detection (TRAP-assay) (Kim et al., 1994; Cairney et al., 2008a). Among the cell lines, four were ALT positive (ALT^+^/TEL^−^), four were telomerase positive (ALT^−^/TEL^+^), and two were ALT/TEL double inactive (ALT^−^/TEL^−^).

Among the liposarcoma samples, nine were ALT positive and eight were telomerase positive. Most of the samples had two technical replicates. In case of multiple microarray probes mapping to the same gene, the probe with highest standard deviation of values was considered. Cell line and tissue data were processed separately. Gene expression values higher than the 0.9 percentile in each of these sets were limited to that percentile value.

Healthy human tissue RNA-seq gene expression data was obtained from the GTEx portal (release V8) in units of transcripts per million (TPM). Fold changes were computed in comparison to the average of non-zero TPM values per gene. Data of the top 15 most common tissues extracted from subjects suffered violent or fast death from natural causes were selected. They were grouped by age and sex. For cross-tissue analysis we selected a maximum number of 10 subjects per age and sex group). For age-dependent analysis of the testis transcriptome all 129 available testis samples were chosen.

### 4.3 Pathway signal flow (PSF) activity and partial influence (PI)

The pathway signal flow (PSF) algorithm (Arakelyan and Nersisyan, 2013; Nersisyan et al., 2016, 2017) was used to asses TMM pathway activity. It computes the activation along the whole pathway based on relative expression values of its member genes and of their interactions. Details are described in the supplement (section Pathway Signal Flow algorithm and Supplementary Figure S1). The PSF algorithm is implemented in the Cytoscape app PSFC (v1.1.8) (Nersisyan et al., 2017). For the specific tasks applied in this work, we have implemented a higher-level app, TMM (v0.8, http://apps.cytoscape.org/apps/tmm). It compares TMM pathway activation patterns with experimental annotations, uses PSFC for pathway activity computation and it also produces reports for TMM phenotype comparison across samples. The app is written in Java (major version 8), the source code is available at https://github.com/lilit-nersisyan/tmm. The app user guide, along with the example datasets and network files can be accessed at the project homepage http://big.sci.am/software/tmm/.

The partial influence (PI) of a source gene estimates the extent to which its expression affects the activity of a downstream target node in the pathway. The PI-value depends on the expression value of the source gene, pathway topology and the expression of other pathway members. It is computed by neutralizing the fold change of the gene to FC= 1, and calculating the log ratio of PSF at the target node before and after neutralizing its expression (Supplementary Figure S4). To compute the mean PI across all the samples in the testis, we have generated a mean sample by averaging fold change values for each gene, and performed PI analyses on it.

## 6 Author Contributions

LN, AA and HB conceived the study. LN and AS performed pathway curation and data analysis. LN designed the software packages and prepared the figures. All the authors contributed to data interpretation and manuscript preparation.

## 8 Acknowledgments

The Genotype-Tissue Expression (GTEx) Project was supported by the Common Fund of the Office of the Director of the National Institutes of Health, and by NCI, NHGRI, NHLBI, NIDA, NIMH, and NINDS. The author(s) acknowledge support from the German Research Foundation (DFG) and Universität Leipzig within the program of Open Access Publishing.

## 10 Tables

## Supplementary data 1

**Table S3.**
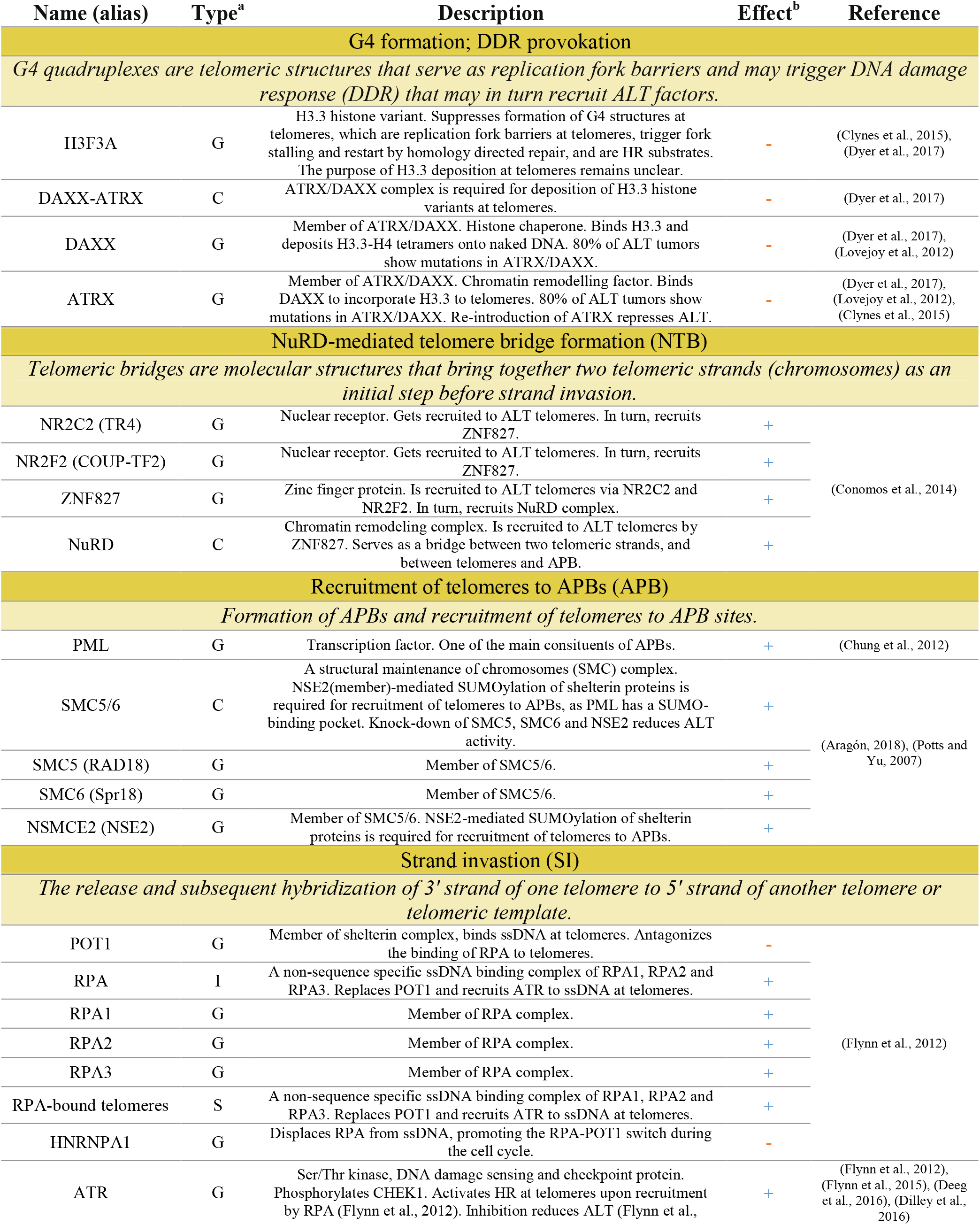
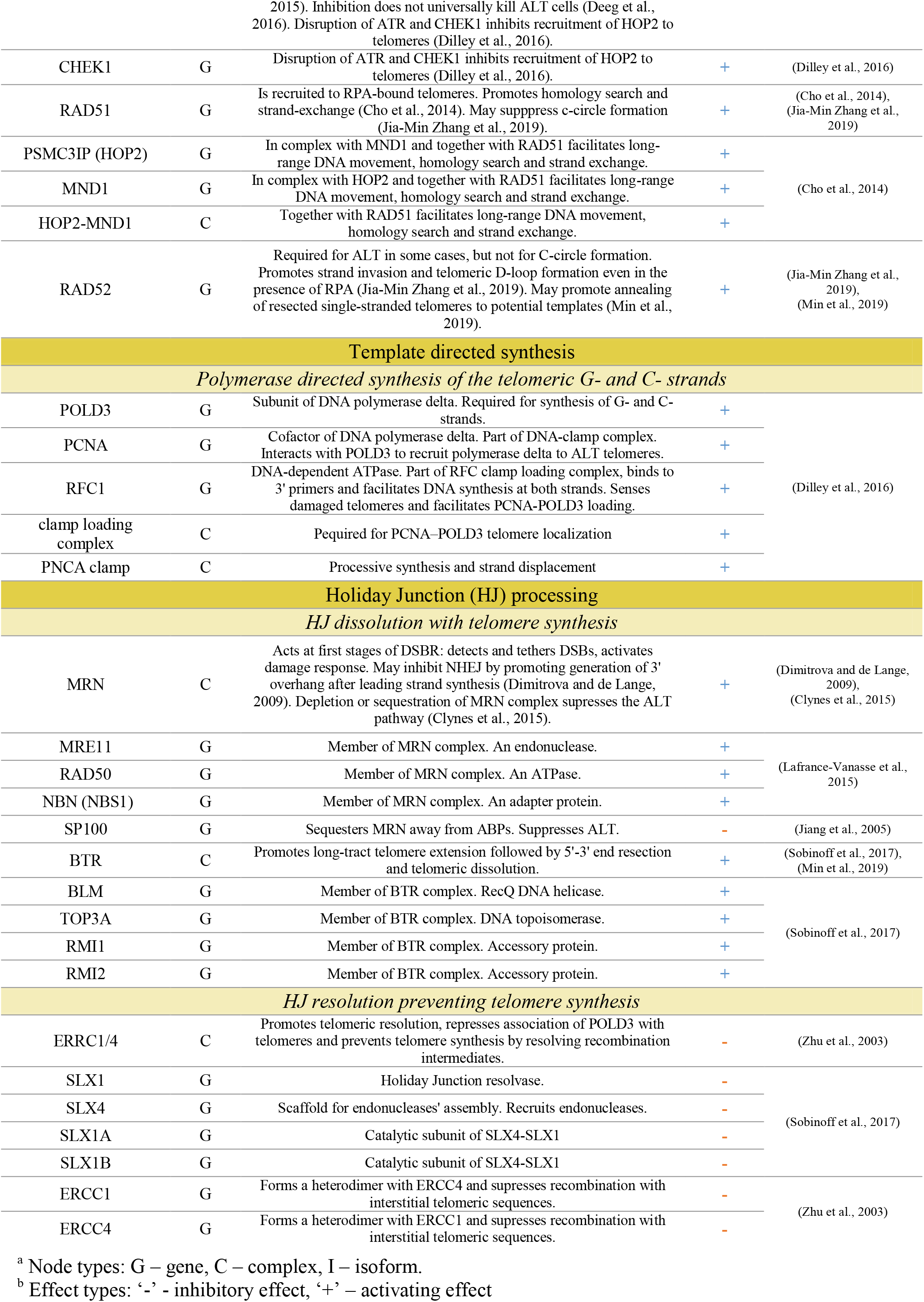
Functional role of the ALT pathway nodes.

**Table S4.**
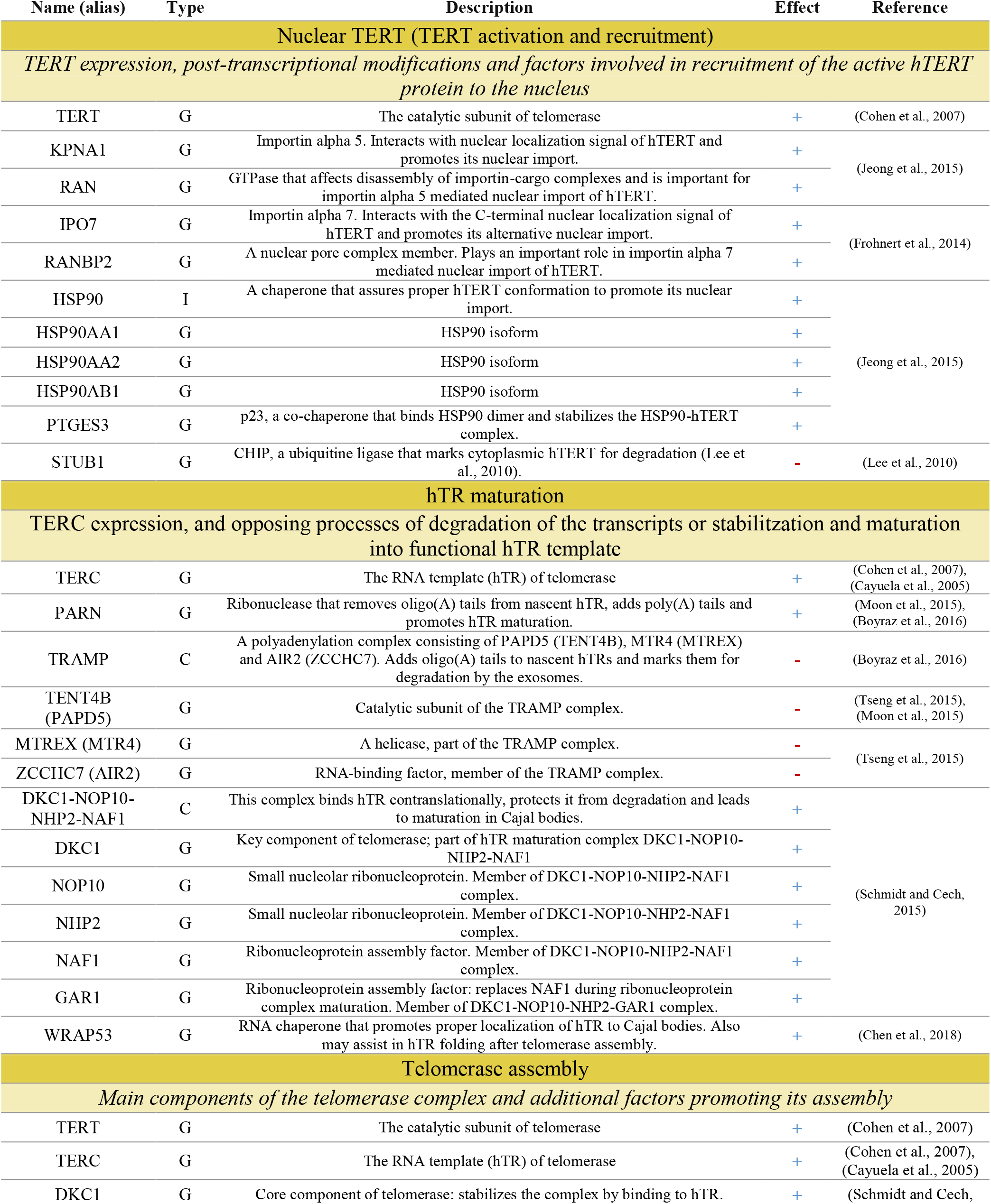
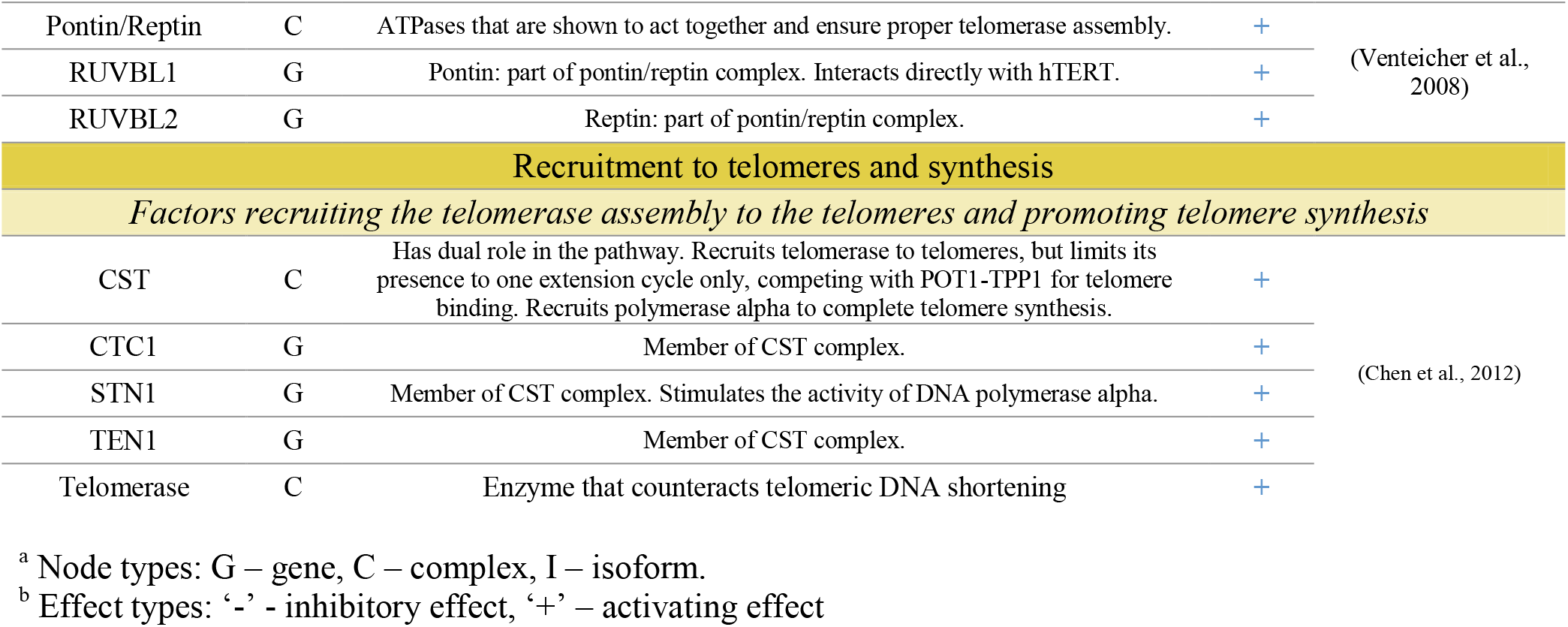
Functional role of the TEL pathway nodes.

### Pathway Signal Flow algorithm

For pathway activity estimation from gene expression data we have used the Pathway Signal Flow (PSF) algorithm, implemented in the *PSFC* app for Cytoscape, v1.1.8 (Nersisyan et al., 2015b). In this particular study, we have made use of a higher level Cytoscape app, *TMM* v0.8 that uses PSFC as a dependency. It compares TMM pathway activation patterns with experimental annotations, uses PSFC for pathway activity computation and it also produces reports for TMM phenotype comparison across samples.The app (optionally) computes the fold change (FC) of each gene by taking the ratio of its expression to the average expression across the samples of the respective data set.

The PSF algorithm (Arakelyan et al., 2013; Nersisyan et al., 2015b, 2016) computes the strength of the signal propagated from the pathway inputs to the outputs through pairwise interactions between nodes, based on their fold change (FC) expression values. In this study, missing values were assigned an FC value of 1. For each source-target interaction, the FC values were multiplied for edges of type *activation* (FC_source_ ∗ FC_target_) and inversely multiplied for edges of type *inhibition* (1/FC_source_ ∗ FC_target_). The signal propagation starts from input nodes, spreads through the intermediate nodes and arrives at the sink nodes (labeled “ALT” and “TEL”, respectively). The PSF scores at the sink nodes reflect the overall activity of the pathways (Figure S1).

The product of multiple signals from many sources was assigned as the signal at the target node (the “multiplication” option in PSFC). PSFC also allows for assigning explicit functions to deal with multiple incoming edges onto specific nodes. We have assigned the function “min” to nodes combining complex subunits (as the gene with minimum expression defines the activity of the complex), and the function “max” to homologous genes, where it is unknown which transcript plays the described role (and assumed that the most expressed one should), and finally we assigned the function “sum” to the linker between two hTERT nuclear import pathways, as each of them independently contribute to the protein‘s entry to the nucleus.

**Figure S1.**
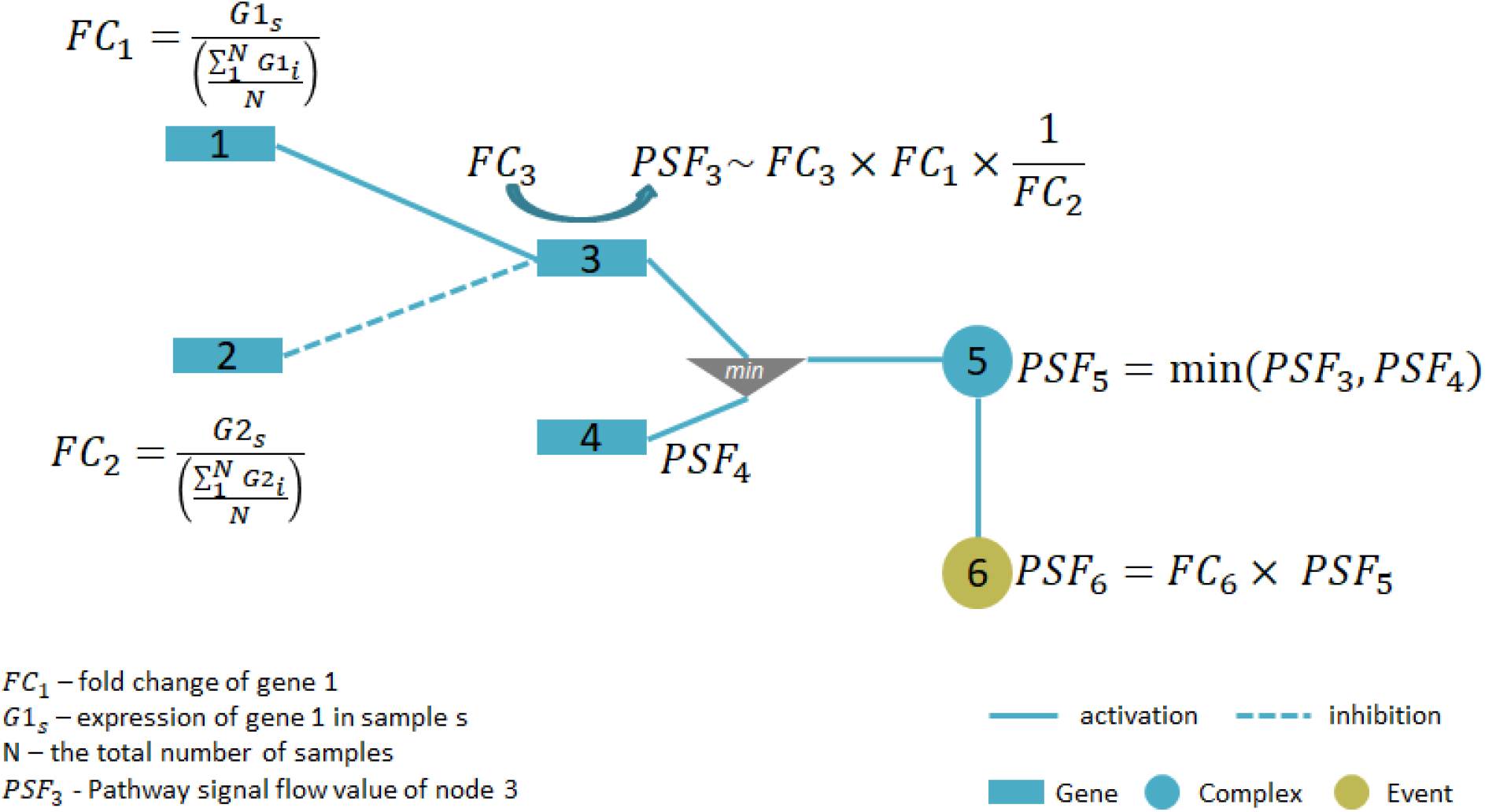
Signal propagation with the pathway signal flow algorithm. The fold change (FC) values for each gene are computed as its expression in the given sample relative to its mean expression across the samples. After initial assignment of FC values, the pathway signal flow (PSF) values are computed by multiplying the source and target values (PSF if already computed or FC if the node doesn‘t have upstream interactors) on the edges of type activation, or performing inverse multiplication on the edges of type inhibition. The operator nodes (min) modulate the PSF values by setting a certain rule on multiple input signals (in this case: the minimum value of the two upstream PSF values on the node of type Complex).

**Figure S2.**
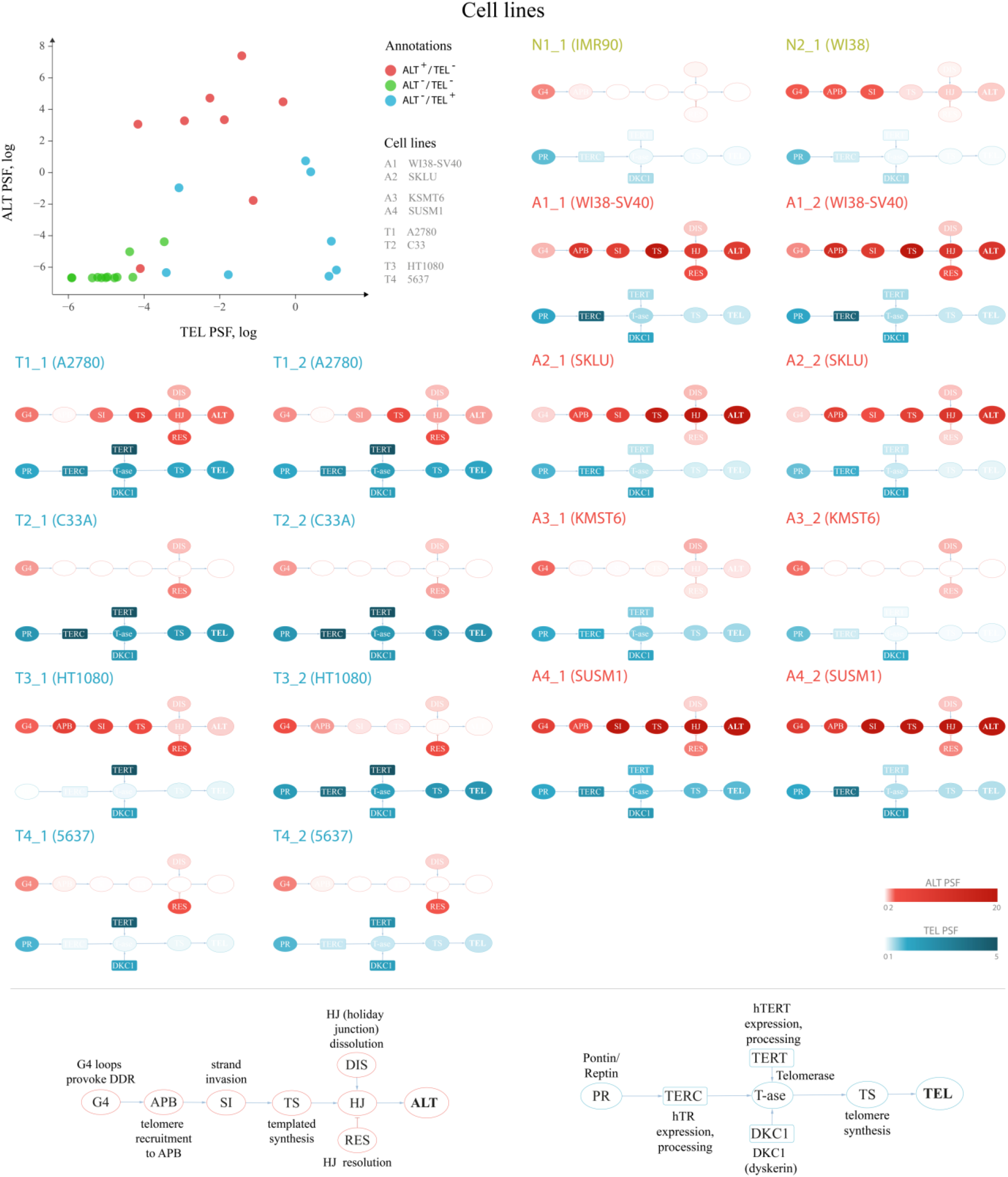
Cell line specific pathway activation patterns at the level of individual pathway branches. Top left: 2D plots with samples placed according to the TEL (x axis) and the ALT (y axis) pathway activity PSF values. The samples are colored according to the experimental TMM annotations. Technical replicates are distinguished with _1 and _2 suffices. Horizontal and vertical dash lines separate ALT^+^ from ALT^−^ and TEL^+^ from TEL^−^ experimentally annotated samples based on support vector machine classification on the ALT and the TEL PSF values. Other panels: Pathway activation patterns at the level of branches in each sample. Bottom: node abbreviations in the pathways.

**Figure S3.**
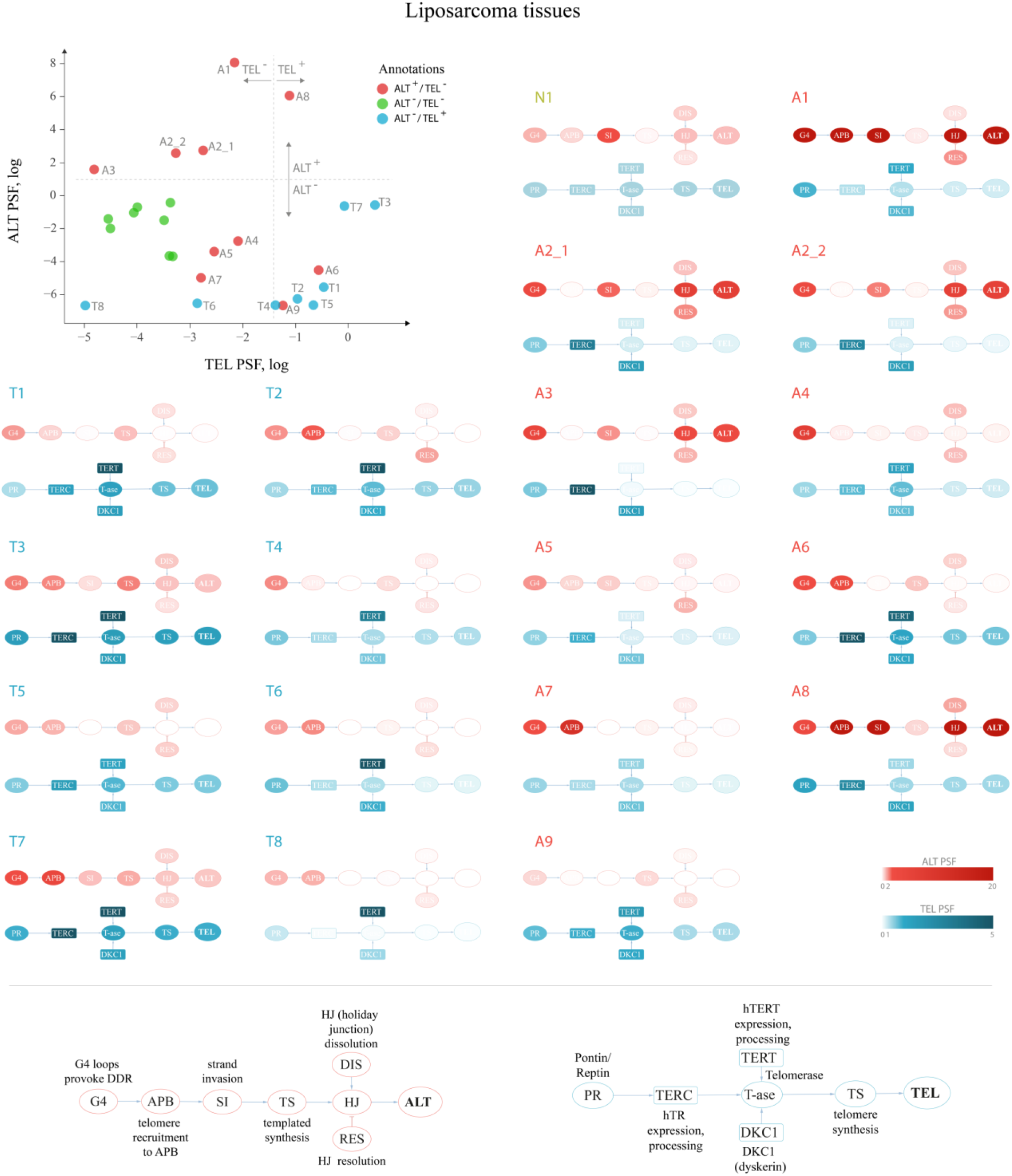
Tissue specific pathway activation patterns at the level of individual pathway branches. Top left: 2D plots with samples placed according to the TEL (x axis) and the ALT (y axis) pathway activity PSF values. The samples are colored according to the experimental TMM annotations. Technical replicates are distinguished with _1 and _2 suffices. Horizontal and vertical dash lines separate ALT^+^ from ALT^−^ and TEL^+^ from TEL^−^ experimentally annotated samples based on support vector machine classification on the ALT and the TEL PSF values. Other panels: Pathway activation patterns at the level of branches in each sample. Bottom: node abbreviations in the pathways.

**Figure S4.**
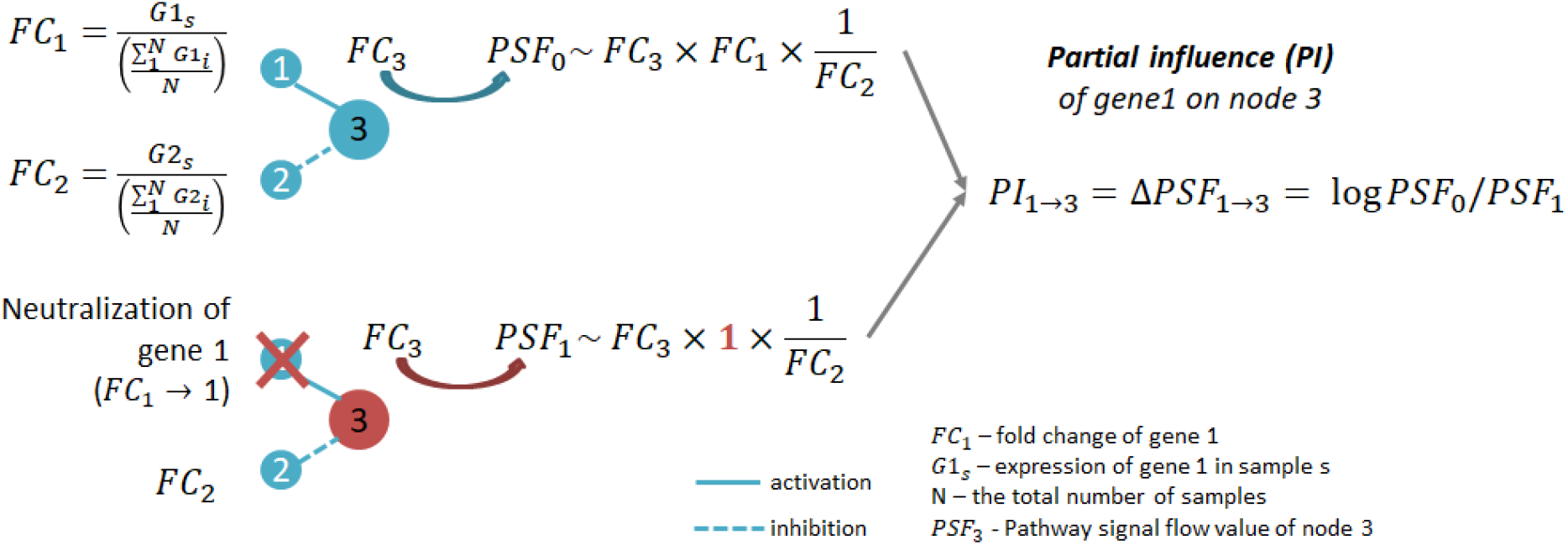
A toy example of computation of the partial influence (PI) of a given source node on the PSF activity of the target node. Top: computation of the PSF activity of the node 3 (PSF_0_), given the fold change (FC) values of the node 1 and 2. Bottom: PSF activity of the node 3 (PSF_1_) after setting the fold change of the node 1 to FC = 1, thus neutralizing its influence. The difference between the PSF activity of the node 3 before and after neutralizing the influence of the node 1 is the partial influence (PI) of the node 1 on node 3 activity.

**Figure S5.**
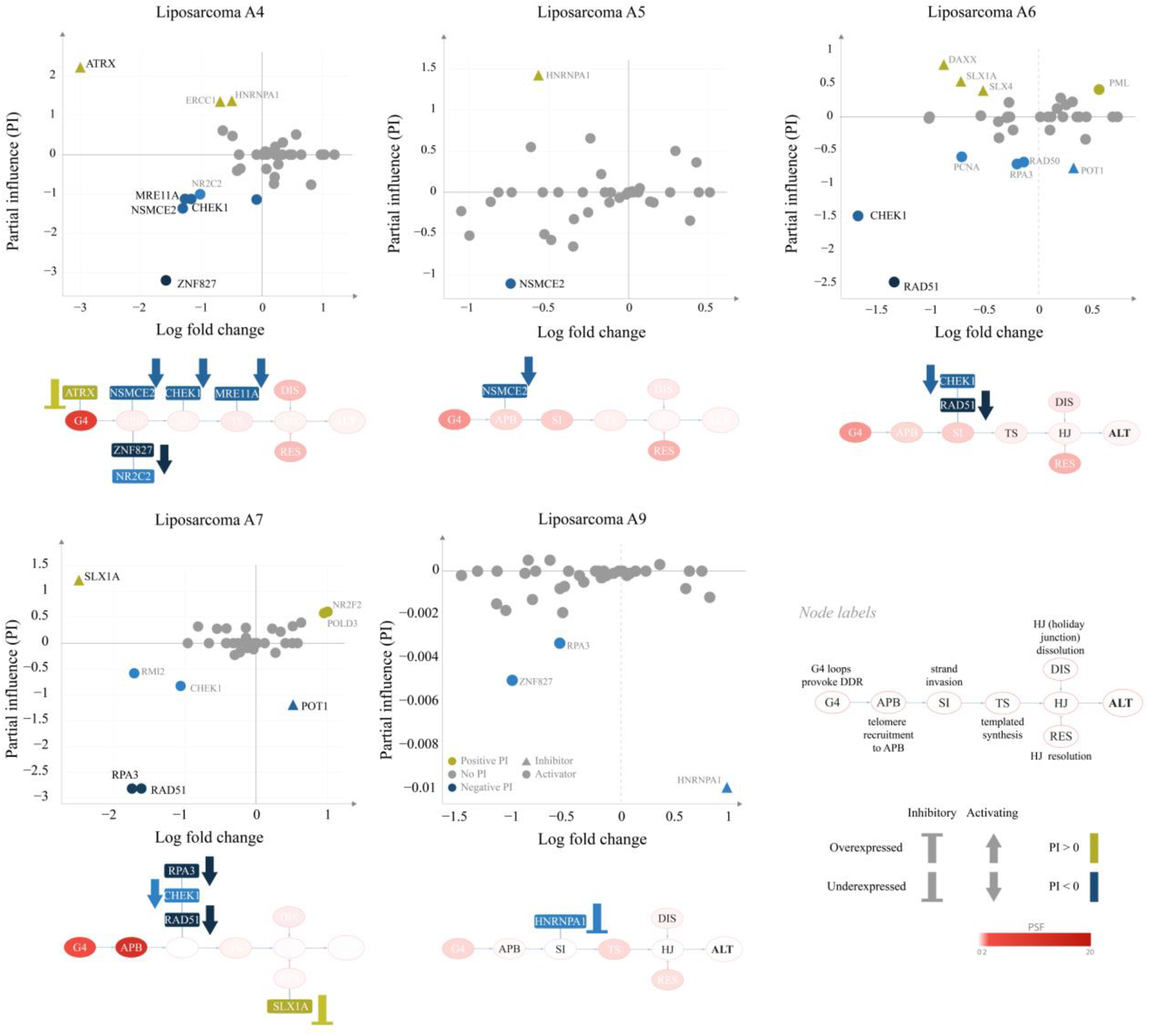
Partial influence of nodes on the ALT pathway activity of the A5 and A9 liposarcoma tissues. Partial influence (PI) of each node is computed as the difference of the TEL node PSF value when the node is set to a fold change (FC) value of 1. The most influential nodes are shown in the context of the pathway branches.

**Figure S6.**
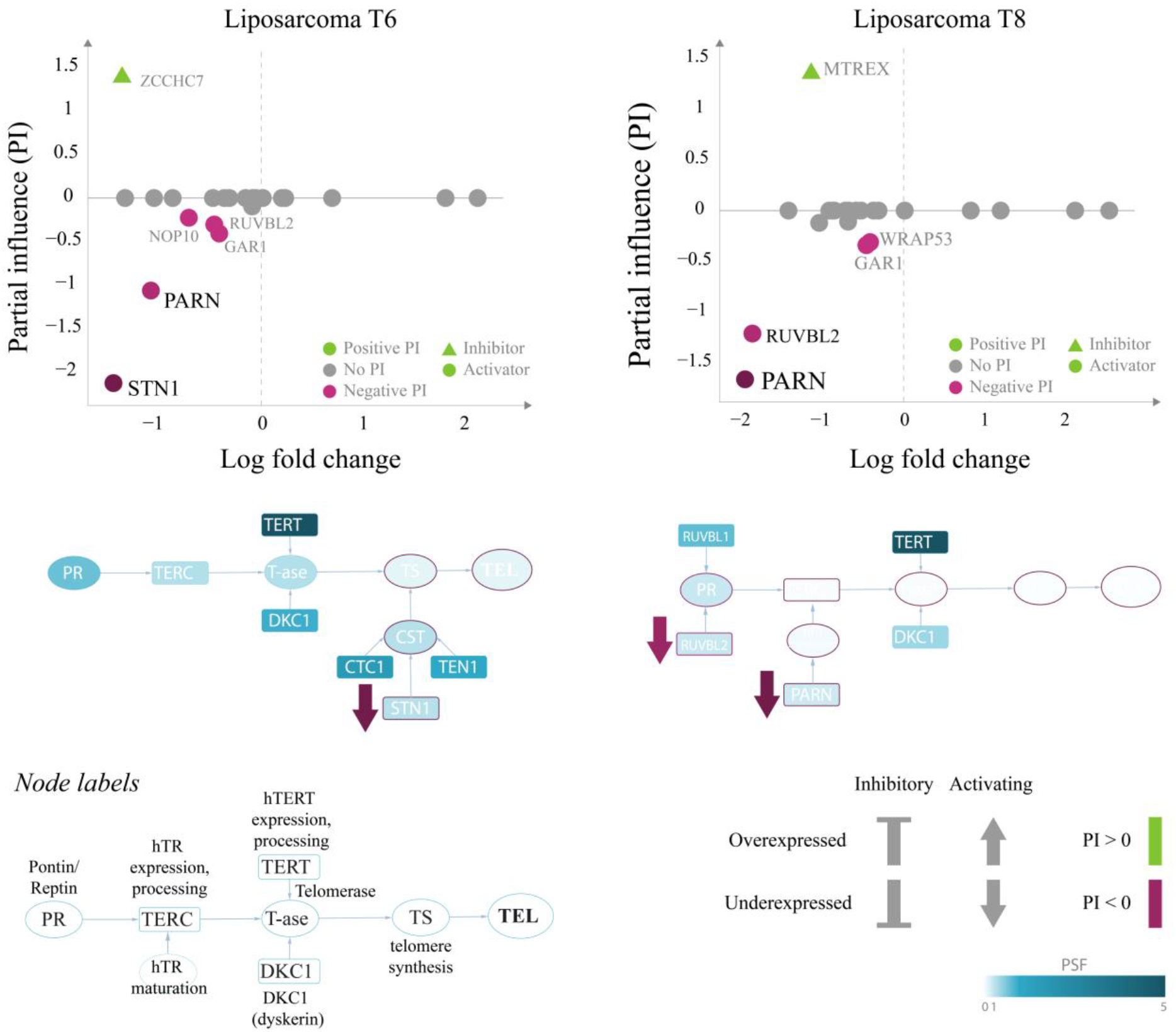
Partial influence of nodes on the TEL pathway activity of the T6 and T8 liposarcoma tissues. Partial influence (PI) of each node is computed as the difference of the TEL node PSF value when the node is set to a fold change (FC) value of 1. The most influential nodes are shown in the context of the pathway branches.

**Table S5.**
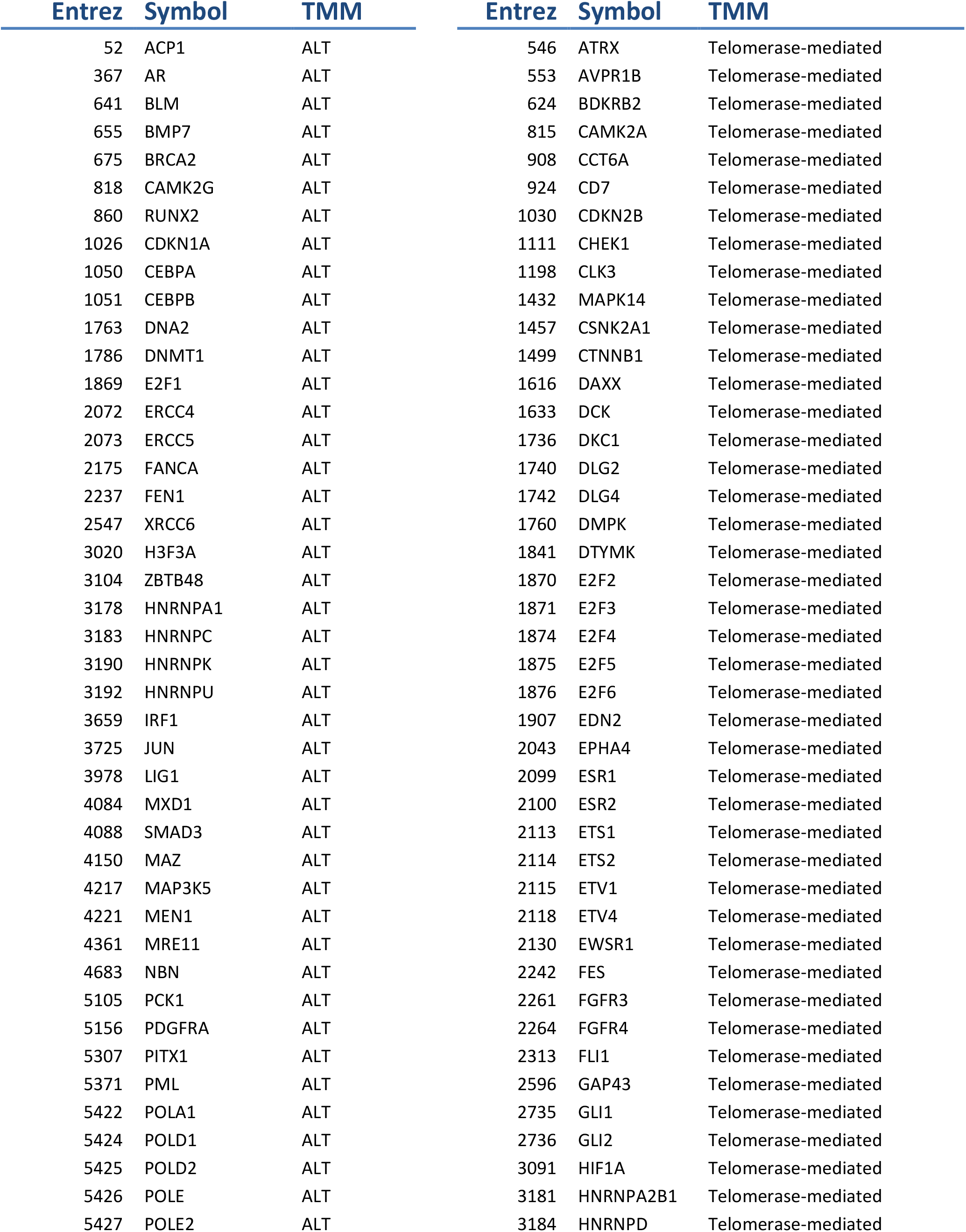
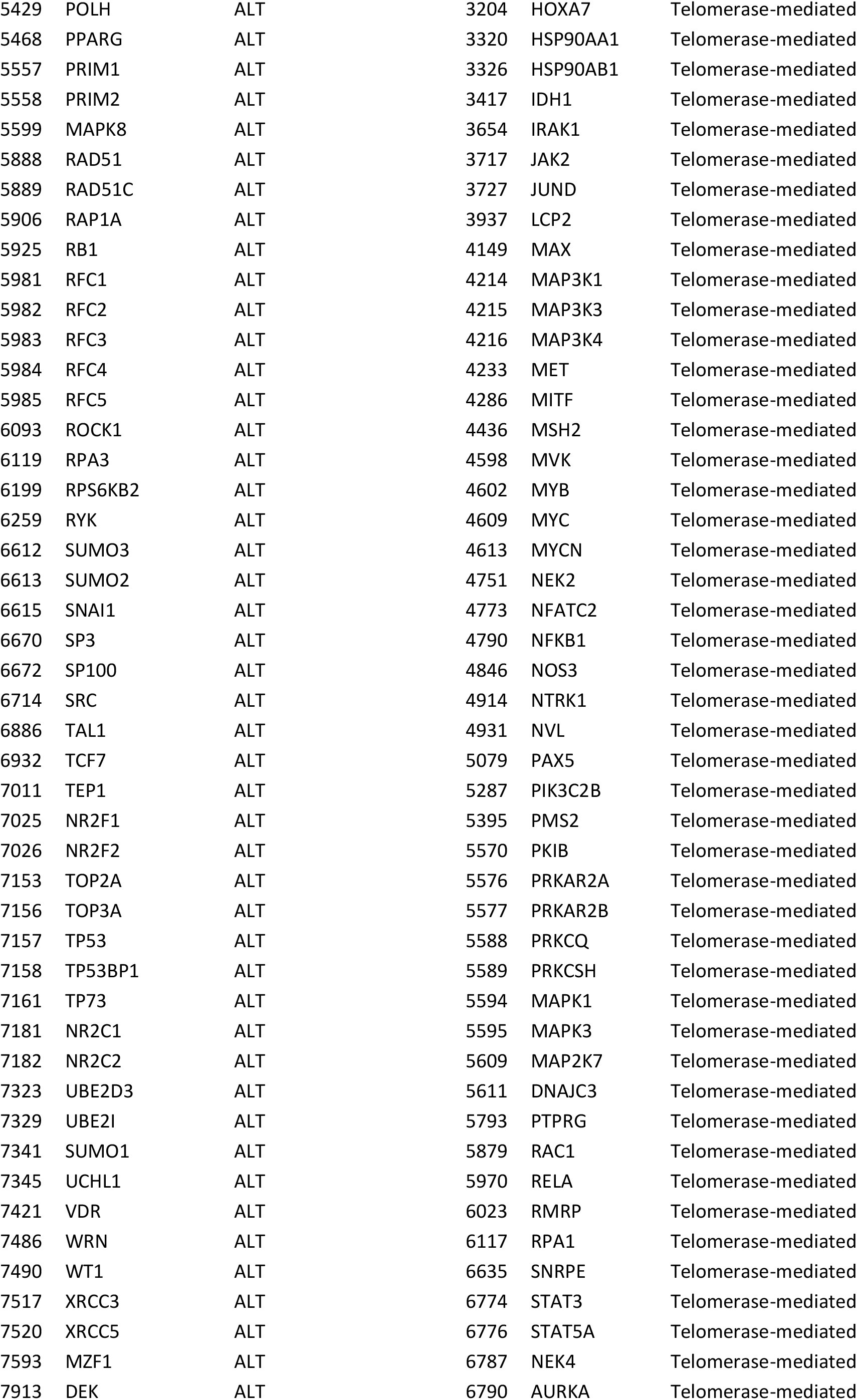
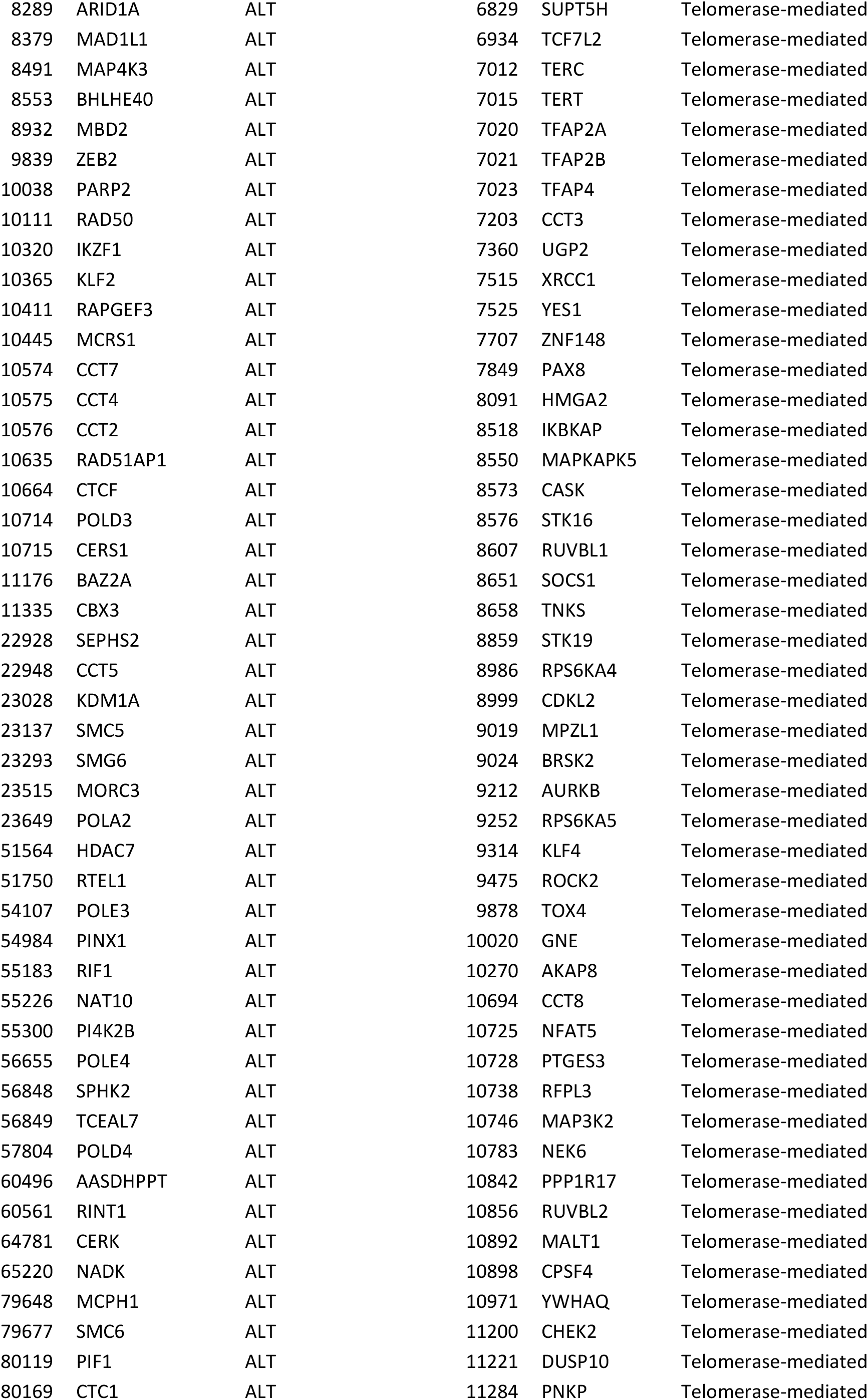
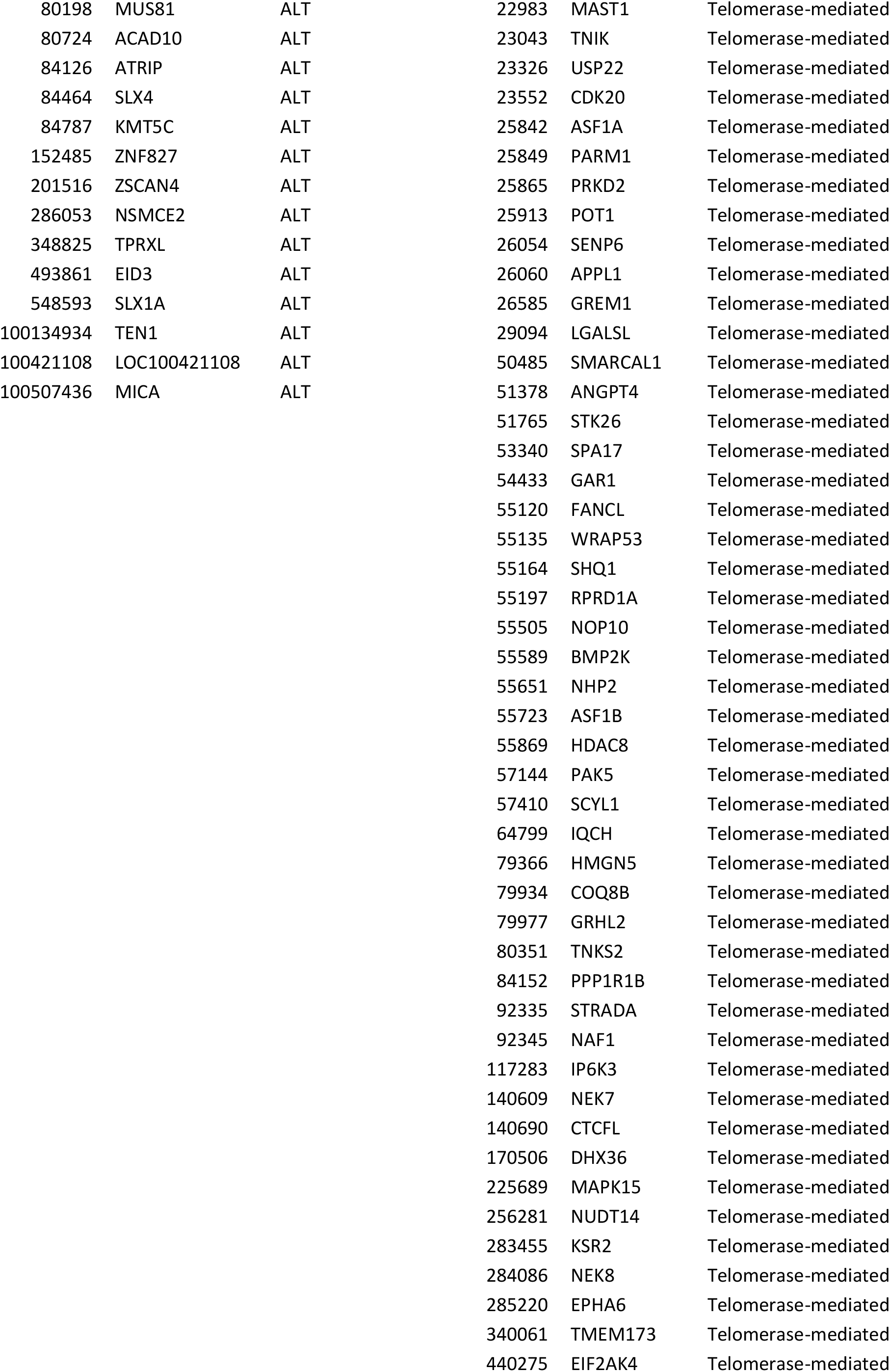
Genes from the TelNet database that are marked as either having a role in ALT or Telomerase-mediated telomere maintenance pathways.

**Figure S7.**
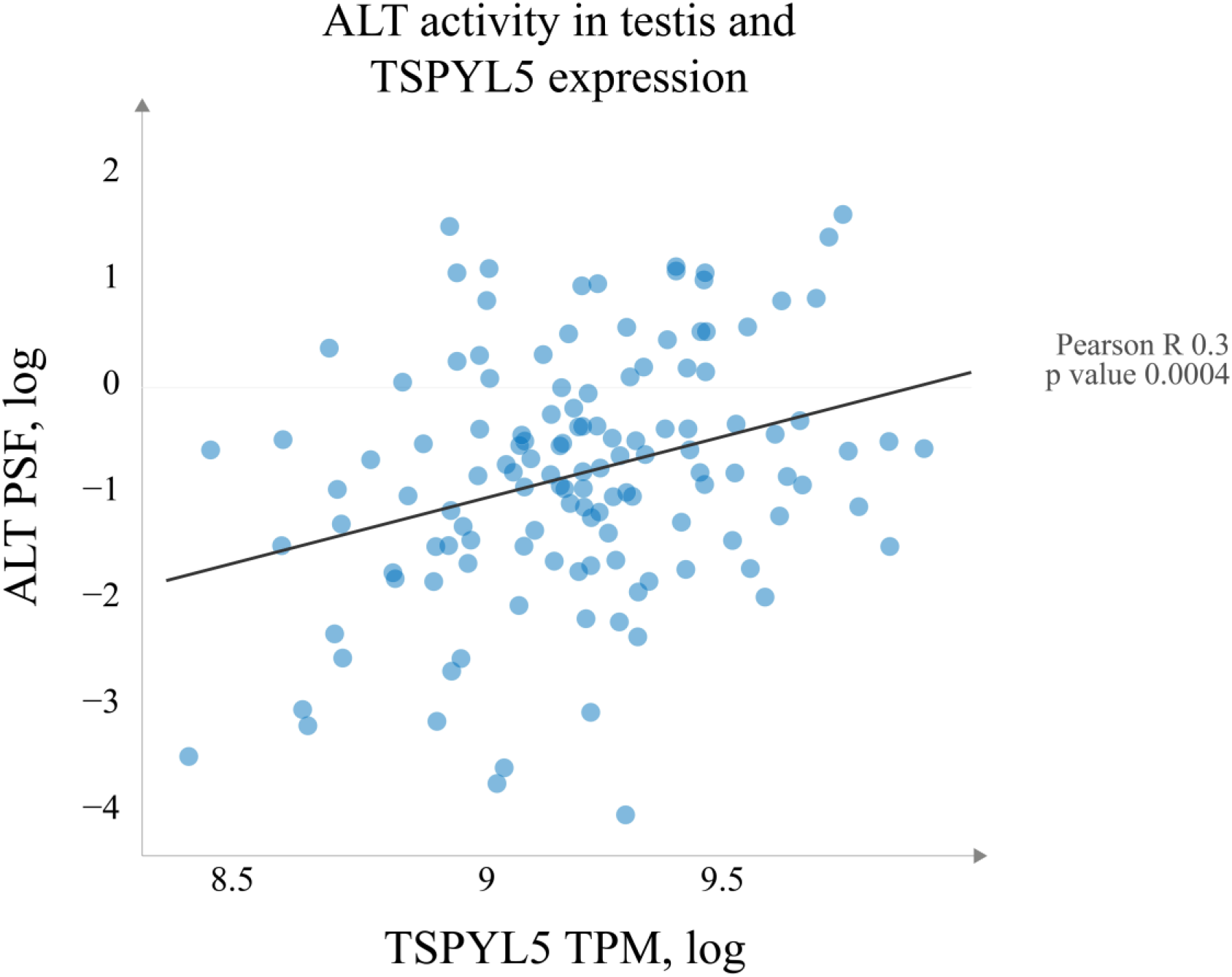
ALT pathway activity and expression of *TSPYL5*. ALT pathway PSF values show slight correlation (Pearson correlation R 0.03) with the expression of the testis-specific Y-encoded-like protein 5 (*TSPYL5*).

## References

Achi, M. V., Ravindranath, N., and Dym, M. (2000). Telomere Length in Male Germ Cells Is Inversely Correlated with Telomerase Activity1. Biol. Reprod. 63, 591–598. doi:10.1095/biolreprod63.2.591.

Alhendi, A. S. N., and Royle, N. J. (2020). The absence of (TCAGGG)n repeats in some telomeres, combined with variable responses to NR2F2 depletion, suggest that this nuclear receptor plays an indirect role in the alternative lengthening of telomeres. Sci. Rep. 10, 20597. doi:10.1038/s41598-020-77606-w.

Aragón, L. (2018). The Smc5/6 Complex: New and Old Functions of the Enigmatic Long-Distance Relative. Annu. Rev. Genet. 52, 89–107. doi:10.1146/annurev-genet-120417-031353.

Arakelyan, A., and Nersisyan, L. (2013). KEGGParser: Parsing and editing KEGG pathway maps in Matlab. Bioinformatics 29, 518–519.

Bechter, O. E., Zou, Y., Walker, W., Wright, W. E., and Shay, J. W. (2004). Telomeric recombination in mismatch repair deficient human colon cancer cells after telomerase inhibition. Cancer Res. 64, 3444–3451. doi:10.1158/0008-5472.CAN-04-0323.

Blackburn, E. H. (1991). Structure and function of telomeres. Nature 350, 569–73. doi:10.1038/350569a0.

Boyraz, B., Moon, D. H., Segal, M., Muosieyiri, M. Z., Aykanat, A., Tai, A. K., et al. (2016). Posttranscriptional manipulation of TERC reverses molecular hallmarks of telomere disease. J. Clin. Invest. 126, 3377–82. doi:10.1172/JCI87547.

Braun, D. M., Chung, I., Kepper, N., Deeg, K. I., and Rippe, K. (2018). TelNet - a database for human and yeast genes involved in telomere maintenance. BMC Genet. 19, 32. doi:10.1186/s12863-018-0617-8.

Cairney, C. J., Hoare, S. F., Daidone, M.-G., Zaffaroni, N., and Keith, W.N. (2008a). High level of telomerase RNA gene expression is associated with chromatin modification, the ALT phenotype and poor prognosis in liposarcoma. Br. J. Cancer 98, 1467–1474. doi:10.1038/sj.bjc.6604328.

Cairney, C. J., Hoare, S. F., Daidone, M. G., Zaffaroni, N., and Keith, W.N. (2008b). High level of telomerase RNA gene expression is associated with chromatin modification, the ALT phenotype and poor prognosis in liposarcoma. Br. J. Cancer 98, 1467–1474. doi:10.1038/sj.bjc.6604328.

Cayuela, M. L., Flores, J. M., and Blasco, M. A. (2005). The telomerase RNA component Terc is required for the tumour-promoting effects of Tert overexpression. EMBO Rep. 6, 268– 274. doi:10.1038/sj.embor.7400359.

Cesare, A. J., and Reddel, R. R. (2008). Telomere uncapping and alternative lengthening of telomeres. Mech. Ageing Dev. 129, 99–108. doi:10.1016/j.mad.2007.11.006.

Chen, L.-Y., Redon, S., and Lingner, J. (2012). The human CST complex is a terminator of telomerase activity. Nature 488, 540–4. doi:10.1038/nature11269.

Chen, L., Roake, C. M., Freund, A., Batista, P. J., Tian, S., Yin, Y. A., et al. (2018). An Activity Switch in Human Telomerase Based on RNA Conformation and Shaped by TCAB1. Cell 174, 218-230.e13. doi:10.1016/j.cell.2018.04.039.

Chen, X., Tang, W., Shi, J. B., Liu, M. M., and Liu, X. (2020). Therapeutic strategies for targeting telomerase in cancer. Med. Res. Rev. 40, 532–585. doi:10.1002/med.21626.

Cho, N. W., Dilley, R. L., Lampson, M. A., and Greenberg, R. A. (2014). Interchromosomal homology searches drive directional ALT telomere movement and synapsis. Cell 159, 108–121. doi:10.1016/j.cell.2014.08.030.

Chow, T. T., Zhao, Y., Mak, S. S., Shay, J. W., and Wright, W. E. (2012). Early and late steps in telomere overhang processing in normal human cells: The position of the final RNA primer drives telomere shortening. Genes Dev. 26, 1167–1178. doi:10.1101/gad.187211.112.

Chung, I., Osterwald, S., Deeg, K. I., and Rippe, K. (2012). PML body meets telomere: the beginning of an ALTernate ending? Nucleus 3, 263–75. doi:10.4161/nucl.20326.

Clynes, D., Jelinska, C., Xella, B., Ayyub, H., Scott, C., Mitson, M., et al. (2015). Suppression of the alternative lengthening of telomere pathway by the chromatin remodelling factor ATRX. Nat. Commun. 6, 7538. doi:10.1038/ncomms8538.

Cohen, S. B., Graham, M. E., Lovrecz, G. O., Bache, N., Robinson, P. J., and Reddel, R. R. (2007). Protein composition of catalytically active human telomerase from immortal cells. Science 315, 1850–3. doi:10.1126/science.1138596.

Conomos, D., Reddel, R. R., and Pickett, H. A. (2014). NuRD-ZNF827 recruitment to telomeres creates a molecular scaffold for homologous recombination. doi:10.1038/nsmb.2877.

Costa, A., Daidone, M. G., Daprai, L., Villa, R., Cantù, S., Pilotti, S., et al. (2006). Telomere maintenance mechanisms in liposarcomas: Association with histologic subtypes and disease progression. Cancer Res. 66, 8918–8924. doi:10.1158/0008-5472.CAN-06-0273.

Dagg, R. A., Pickett, H. A., Neumann, A. A., Napier, C. E., Henson, J. D., Teber, E. T., et al. (2017). Extensive Proliferation of Human Cancer Cells with Ever-Shorter Telomeres. Cell Rep. 19, 2544–2556. doi:10.1016/j.celrep.2017.05.087.

Deeg, K. I., Chung, I., Bauer, C., and Rippe, K. (2016). Cancer Cells with Alternative Lengthening of Telomeres Do Not Display a General Hypersensitivity to ATR Inhibition. Front. Oncol. 6, 186. doi:10.3389/fonc.2016.00186.

Demanelis, K., Jasmine, F., Chen, L. S., Chernoff, M., Tong, L., Delgado, D., et al. (2020). Determinants of telomere length across human tissues. Science (80-.). 369. doi:10.1126/SCIENCE.AAZ6876.

Dilley, R. L., Verma, P., Cho, N. W., Winters, H. D., Wondisford, A. R., and Greenberg, R. A. (2016). Break-induced telomere synthesis underlies alternative telomere maintenance. Nature 539, 54–58. doi:10.1038/nature20099.

Dimitrova, N., and de Lange, T. (2009). Cell Cycle-Dependent Role of MRN at Dysfunctional Telomeres: ATM Signaling-Dependent Induction of Nonhomologous End Joining (NHEJ) in G1 and Resection-Mediated Inhibition of NHEJ in G2. Mol. Cell. Biol. 29, 5552–5563. doi:10.1128/mcb.00476-09.

Dyer, M. A., Qadeer, Z. A., Valle-Garcia, D., and Bernstein, E. (2017). ATRX and DAXX: Mechanisms and Mutations. Cold Spring Harb. Perspect. Med. 7, a026567. doi:10.1101/cshperspect.a026567.

Episkopou, H., Diman, A., Claude, E., Viceconte, N., and Decottignies, A. (2019). TSPYL5 Depletion Induces Specific Death of ALT Cells through USP7-Dependent Proteasomal Degradation of POT1. Mol. Cell 75, 469-482.e6. doi:10.1016/j.molcel.2019.05.027.

Fajkus, J. (2006). Detection of telomerase activity by the TRAP assay and its variants and alternatives. Clin. Chim. Acta 371, 25–31. doi:10.1016/j.cca.2006.02.039.

Flynn, R. L., Chang, S., and Zou, L. (2012). RPA and POT1: friends or foes at telomeres? Cell Cycle 11, 652–7. doi:10.4161/cc.11.4.19061.

Flynn, R. L., Cox, K. E., Jeitany, M., Wakimoto, H., Bryll, A. R., Ganem, N. J., et al. (2015). Alternative lengthening of telomeres renders cancer cells hypersensitive to ATR inhibitors. Science 347, 273–7. doi:10.1126/science.1257216.

Frohnert, C., Hutten, S., Wälde, S., Nath, A., and Kehlenbach, R. H. (2014). Importin 7 and Nup358 promote nuclear import of the protein component of human telomerase. PLoS One 9. doi:10.1371/journal.pone.0088887.

Gan, Y., Mo, Y., Johnston, J., Lu, J., Wientjes, M. G., and Au, J. L.-S. (2002). Telomere maintenance in telomerase-positive human ovarian SKOV-3 cells cannot be retarded by complete inhibition of telomerase. FEBS Lett. 527, 10–14. doi:10.1016/S0014-5793(02)03141-1.

Gocha, A. R. S., Nuovo, G., Iwenofu, O. H., and Groden, J. (2013). Human sarcomas are mosaic for telomerase-dependent and telomerase-independent telomere maintenance mechanisms: Implications for telomere-based therapies. Am. J. Pathol. 182, 41–48. doi:10.1016/j.ajpath.2012.10.001.

Greenberg, R. a (2005). Telomeres, crisis and cancer. Curr. Mol. Med. 5, 213–8. doi:10.2174/1566524053586590.

Hakobyan, A., Nersisyan, L., and Arakelyan, A. (2016). Quantitative trait association study for mean telomere length in the South Asian genomes. Bioinformatics 32, 1697–700. doi:10.1093/bioinformatics/btw027.

Henson, J. D., Neumann, A. A., Yeager, T. R., and Reddel, R. R. (2002). Alternative lengthening of telomeres in mammalian cells. Oncogene 21, 598–610. doi:10.1038/sj.onc.1205058.

Henson, J. D., and Reddel, R. R. (2010). Assaying and investigating Alternative Lengthening of Telomeres activity in human cells and cancers. FEBS Lett. 584, 3800–3811. doi:10.1016/j.febslet.2010.06.009.

Hug, N., and Lingner, J. (2006). Telomere length homeostasis. Chromosoma 115, 413–425. doi:10.1007/s00412-006-0067-3.

Jeong, S. A., Kim, K., Lee, J. H., Cha, J. S., Khadka, P., Cho, H. S., et al. (2015). Akt-mediated phosphorylation increases the binding affinity of hTERT for importin α to promote nuclear translocation. J. Cell Sci. doi:10.1242/jcs.166132.

Jia-Min Zhang, A., Yadav, T., Ouyang, J., Lan, L., and Zou Correspondence, L. (2019). Alternative Lengthening of Telomeres through Two Distinct Break-Induced Replication Pathways. CellReports 26, 955-968.e3. doi:10.1016/j.celrep.2018.12.102.

Jiang, W.-Q., Zhong, Z.-H., Henson, J. D., Neumann, A. A., Chang, A. C.-M., and Reddel, R. R. (2005). Suppression of Alternative Lengthening of Telomeres by Sp100-Mediated Sequestration of the MRE11/RAD50/NBS1 Complex. Mol. Cell. Biol. 25, 2708–2721. doi:10.1128/MCB.25.7.2708-2721.2005.

Kim, N. W., Piatyszek, M. A., Prowse, K. R., Harley, C. B., West, M. D., Ho, P. L. C., et al. (1994). Specific association of human telomerase activity with immortal cells and cancer. Science (80-.). 266, 2011–2015. doi:10.1126/science.7605428.

Lafferty-Whyte, K., Cairney, C. J., Will, M. B., Serakinci, N., Daidone, M.-G., Zaffaroni, N., et al. (2009). A gene expression signature classifying telomerase and ALT immortalization reveals an hTERT regulatory network and suggests a mesenchymal stem cell origin for ALT. Oncogene 28, 3765–74. doi:10.1038/onc.2009.238.

Lafrance-Vanasse, J., Williams, G. J., and Tainer, J. A. (2015). Envisioning the dynamics and flexibility of Mre11-Rad50-Nbs1 complex to decipher its roles in DNA replication and repair. Prog. Biophys. Mol. Biol. 117, 182–193. doi:10.1016/j.pbiomolbio.2014.12.004.

Lee, J. H., Khadka, P., Baek, S. H., and Chung, I. K. (2010). CHIP promotes hTERT degradation and negatively regulates telomerase activity. J. Biol. Chem. 285, 42033–42045. doi:10.1074/jbc.M110.149831.

Liu, L., Bailey, S. M., Okuka, M., Muñoz, P., Li, C., Zhou, L., et al. (2007). Telomere lengthening early in development. Nat. Cell Biol. 9, 1436–1441. doi:10.1038/ncb1664.

Lovejoy, C. A., Li, W., Reisenweber, S., Thongthip, S., Bruno, J., de Lange, T., et al. (2012). Loss of ATRX, genome instability, and an altered DNA damage response are hallmarks of the alternative lengthening of telomeres pathway. PLoS Genet. 8, e1002772. doi:10.1371/journal.pgen.1002772.

Martĺnez, P., and Blasco, M. A. (2015). Replicating through telomeres: a means to an end. Trends Biochem. Sci. 40, 504–15. doi:10.1016/j.tibs.2015.06.003.

Min, J., Wright, W. E., and Shay, J. W. (2019). Clustered telomeres in phase-separated nuclear condensates engage mitotic DNA synthesis through BLM and RAD52. Genes Dev. 33, 814–827. doi:10.1101/gad.324905.119.

Moon, D. H., Segal, M., Boyraz, B., Guinan, E., Hofmann, I., Cahan, P., et al. (2015). Poly(A)-specific ribonuclease (PARN) mediates 3′-end maturation of the telomerase RNA component. Nat. Genet. 47, 1482–1488. doi:10.1038/ng.3423.

Nersisyan, L. (2017). Telomere Analysis Based on High-Throughput Multi-Omics Data. Available at: urn:nbn:de:bsz:15-qucosa2-162974.

Nersisyan, L., Hopp, L., Loeffler-Wirth, H., Galle, J., Loeffler, M., Arakelyan, A., et al. (2019). Telomere Length Maintenance and Its Transcriptional Regulation in Lynch Syndrome and Sporadic Colorectal Carcinoma. Front. Oncol. 9.

Nersisyan, L., Johnson, G., Riel-Mehan, M., Pico, A. R., and Arakelyan, A. (2017). PSFC: a Pathway Signal Flow Calculator App for Cytoscape [version 2; peer review: 2 approved]. F1000Research 4. doi:10.12688/f1000research.6706.2.

Nersisyan, L., Löffler-Wirth, H., Arakelyan, A., and Binder, H. (2016). Gene Set-and Pathway-Centered Knowledge Discovery Assigns Transcriptional Activation Patterns in Brain, Blood, and Colon Cancer: A Bioinformatics Perspective. Int. J. Knowl. Discov. Bioinforma. 4, 24. doi:10.4018/IJKDB.2014070104.

Neumann, A. A., Watson, C. M., Noble, J. R., Pickett, H. A., Tam, P. P. L., and Reddel, R. R. (2013). Alternative lengthening of telomeres in normal mammalian somatic cells. Genes Dev. 27, 18–23. doi:10.1101/gad.205062.112.

Novakovic, B., Napier, C. E., Vryer, R., Dimitriadis, E., Manuelpillai, U., Sharkey, A., et al. (2016). DNA methylation mediated up-regulation of TERRA non-coding RNA is coincident with elongated telomeres in the human placenta. Mol. Hum. Reprod. 22, 791–799. doi:10.1093/molehr/gaw053.

Ozturk, S. (2015). Telomerase activity and telomere length in male germ cells. Biol. Reprod. 92, 53–54. doi:10.1095/biolreprod.114.124008.

Paradis, V., Dargère, D., Laurendeau, I., Benoît, G., Vidaud, M., Jardin, A., et al. (1999). Expression of the RNA component of human telomerase (hTR) in prostate cancer, prostatic intraepithelial neoplasia, and normal prostate tissue. J. Pathol. 189, 213–218. doi:10.1002/(SICI)1096-9896(199910)189:2<213::AID-PATH417>3.0.CO;2-A.

Pickett, H. A., and Reddel, R. R. (2015). Molecular mechanisms of activity and derepression of alternative lengthening of telomeres. Nat. Struct. Mol. Biol. 22, 875–880. doi:10.1038/nsmb.3106.

Potts, P. R., and Yu, H. (2007). The SMC5/6 complex maintains telomere length in ALT cancer cells through SUMOylation of telomere-binding proteins. Nat. Struct. Mol. Biol. 14, 581–90. doi:10.1038/nsmb1259.

Recagni, M., Bidzinska, J., Zaffaroni, N., and Folini, M. (2020). The Role of Alternative Lengthening of Telomeres Mechanism in Cancer: Translational and Therapeutic Implications. Cancers (Basel). 12, 949. doi:10.3390/cancers12040949.

Rice, C., and Skordalakes, E. (2016). Structure and function of the telomeric CST complex. Comput. Struct. Biotechnol. J. 14, 161–7. doi:10.1016/j.csbj.2016.04.002.

Schmidt, J. C., and Cech, T. R. (2015). Human telomerase: Biogenesis, trafficking, recruitment, and activation. Genes Dev. 29, 1095–1105. doi:10.1101/gad.263863.115.

Shay, J. W. (2016). Role of Telomeres and Telomerase in Aging and Cancer. Cancer Discov. 6, 584–593. doi:10.1158/2159-8290.CD-16-0062.

Shay, J. W., Reddel, R. R., and Wright, W. E. (2012). Cancer: Cancer and telomeres - An alternative to telomerase. Science (80-.). doi:10.1126/science.1222394.

Sobinoff, A. P., Allen, J. A., Neumann, A. A., Yang, S. F., Walsh, M. E., Henson, J. D., et al. (2017). BLM and SLX4 play opposing roles in recombination-dependent replication at human telomeres. EMBO J. 36, 2907–2919. doi:10.15252/embj.201796889.

Sobinoff, A. P., and Pickett, H. A. (2017). Alternative Lengthening of Telomeres: DNA Repair Pathways Converge. Trends Genet. 33, 921–932. doi:10.1016/J.TIG.2017.09.003.

Sobinoff, A. P., and Pickett, H. A. (2020). Mechanisms that drive telomere maintenance and recombination in human cancers. Curr. Opin. Genet. Dev. 60, 25–30. doi:10.1016/j.gde.2020.02.006.

Sugarman, E. T., Zhang, G., and Shay, J. W. (2019). In perspective: An update on telomere targeting in cancer. Mol. Carcinog. 58, 1581–1588. doi:10.1002/mc.23035.

Tseng, C.-K., Wang, H.-F., Burns, A. M., Schroeder, M. R., Gaspari, M., and Baumann, P. (2015). Human Telomerase RNA Processing and Quality Control. Cell Rep. 13, 2232–2243. doi:10.1016/j.celrep.2015.10.075.

Venteicher, A. S., Meng, Z., Mason, P. J., Veenstra, T. D., and Artandi, S. E. (2008). Identification of ATPases Pontin and Reptin as Telomerase Components Essential for Holoenzyme Assembly. Cell 132, 945–957. doi:10.1016/j.cell.2008.01.019.

Verma, P., Dilley, R. L., Gyparaki, M. T., and Greenberg, R. A. (2018). “Direct Quantitative Monitoring of Homology-Directed DNA Repair of Damaged Telomeres,” in Methods in Enzymology doi:10.1016/bs.mie.2017.11.010.

Villa, R., Daidone, M. G., Motta, R., Venturini, L., De Marco, C., Vannelli, A., et al. (2008). Multiple mechanisms of telomere maintenance exist and differentially affect clinical outcome in diffuse malignant peritoneal mesothelioma. Clin. Cancer Res. 14, 4134–4140. doi:10.1158/1078-0432.CCR-08-0099.

Yuan, X., Larsson, C., and Xu, D. (2019). Mechanisms underlying the activation of TERT transcription and telomerase activity in human cancer: old actors and new players. Oncogene 38, 6172–6183. doi:10.1038/s41388-019-0872-9.

Zhang, J.-M., Genois, M.-M., Ouyang, J., Lan, L., and Zou, L. (2021). Alternative lengthening of telomeres is a self-perpetuating process in ALT-associated PML bodies. Mol. Cell. doi:10.1016/j.molcel.2020.12.030.

Zhu, X. D., Niedernhofer, L., Kuster, B., Mann, M., Hoeijmakers, J. H. J., and De Lange, T. (2003). ERCC1/XPF Removes the 3′ Overhang from Uncapped Telomeres and Represses Formation of Telomeric DNA-Containing Double Minute Chromosomes. Mol. Cell. doi:10.1016/S1097-2765(03)00478-7.

## References

Aragón, L. (2018). The Smc5/6 Complex: New and Old Functions of the Enigmatic Long-Distance Relative. Annu. Rev. Genet. 52, 89– 107. doi:10.1146/annurev-genet-120417-031353.

Cayuela, M. L., Flores, J. M., and Blasco, M. A. (2005). The telomerase RNA component Terc is required for the tumour-promoting effects of Tert overexpression. EMBO Rep. 6, 268–274. doi:10.1038/sj.embor.7400359.

Chen, L., Roake, C. M., Freund, A., Batista, P. J., Tian, S., Yin, Y. A., et al. (2018). An Activity Switch in Human Telomerase Based on RNA Conformation and Shaped by TCAB1. Cell 174, 218–230.e13. doi:10.1016/j.cell.2018.04.039.

Jia-Min Zhang, A., Yadav, T., Ouyang, J., Lan, L., and Zou Correspondence, L. (2019). Alternative Lengthening of Telomeres through Two Distinct Break-Induced Replication Pathways. CellReports 26, 955–968.e3. doi:10.1016/j.celrep.2018.12.102.

Jiang, W.-Q., Zhong, Z.-H., Henson, J. D., Neumann, A. A., Chang, A. C.-M., and Reddel, R. R. (2005). Suppression of Alternative Lengthening of Telomeres by Sp100-Mediated Sequestration of the MRE11/RAD50/NBS1 Complex. Mol. Cell. Biol. 25, 2708– 2721. doi:10.1128/MCB.25.7.2708-2721.2005.

