## Supplementary data 2 for "Telomere maintenance pathway activity analysis enables tissue- and gene-level inferences": supplementary_data_2.html

Supplementary data 1


### Supplementary data 1

###### Lilit Nersisyan

- Correlation heatmaps
  - Dataset 1: cell lines
  - Dataset 2: liposarcoma tissues and hMSCs
  - Pairwise correlation between technical replicates
    - Dataset 1: cell lines
    - Dataset 2: liposarcoma tissues and hMSCs

The plots below show the correlation of gene expression between biological and technical replicates of the analyzed samples deriving from two datasets. All the genes or those belonging to the TEL and ALT pathways reconstructed by us are analyzed.

The sample names are followed by \_1 or \_2, indicating the first and the second technical replicates.

Tags:

normal - mortal cell lines or human mesenchymal stem cells (hMSC)

ALT - samples with high amount of APB bodies

telomerase - samples positive on telomerase TRAP assay

### Correlation heatmaps

Two heatmaps for each gene set are drawn: a not clustered one followd by a clustered one. No scaling of gene expression values has been applied.

##### Dataset 1: cell lines

##### Dataset 2: liposarcoma tissues and hMSCs

#### Pairwise correlation between technical replicates

##### Dataset 1: cell lines

##### Dataset 2: liposarcoma tissues and hMSCs
