## Supplementary figures and images for "Telomere maintenance pathway activity analysis enables tissue- and gene-level inferences"

### ALT_5637_1.jpg

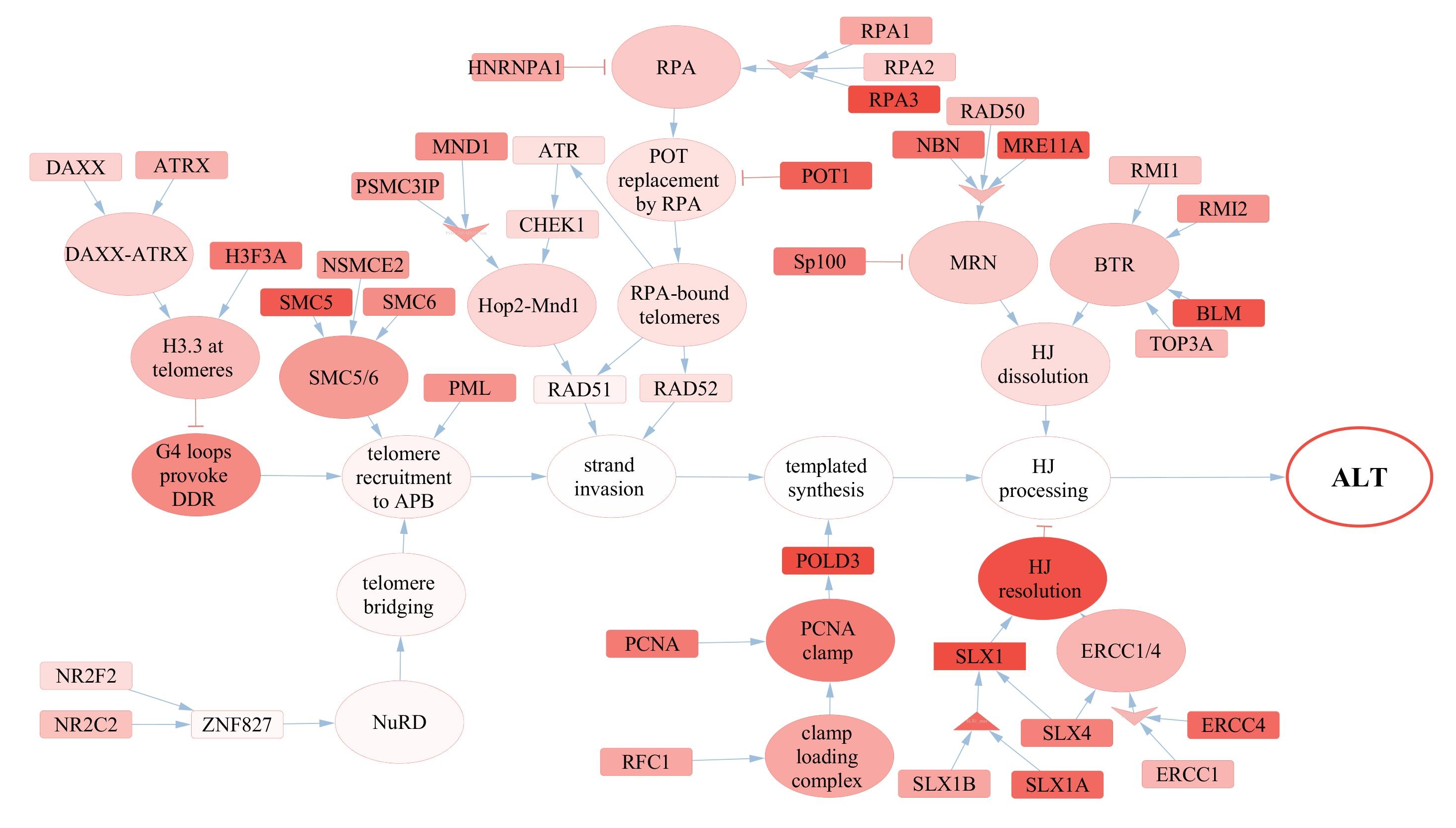

### ALT_A1.jpg

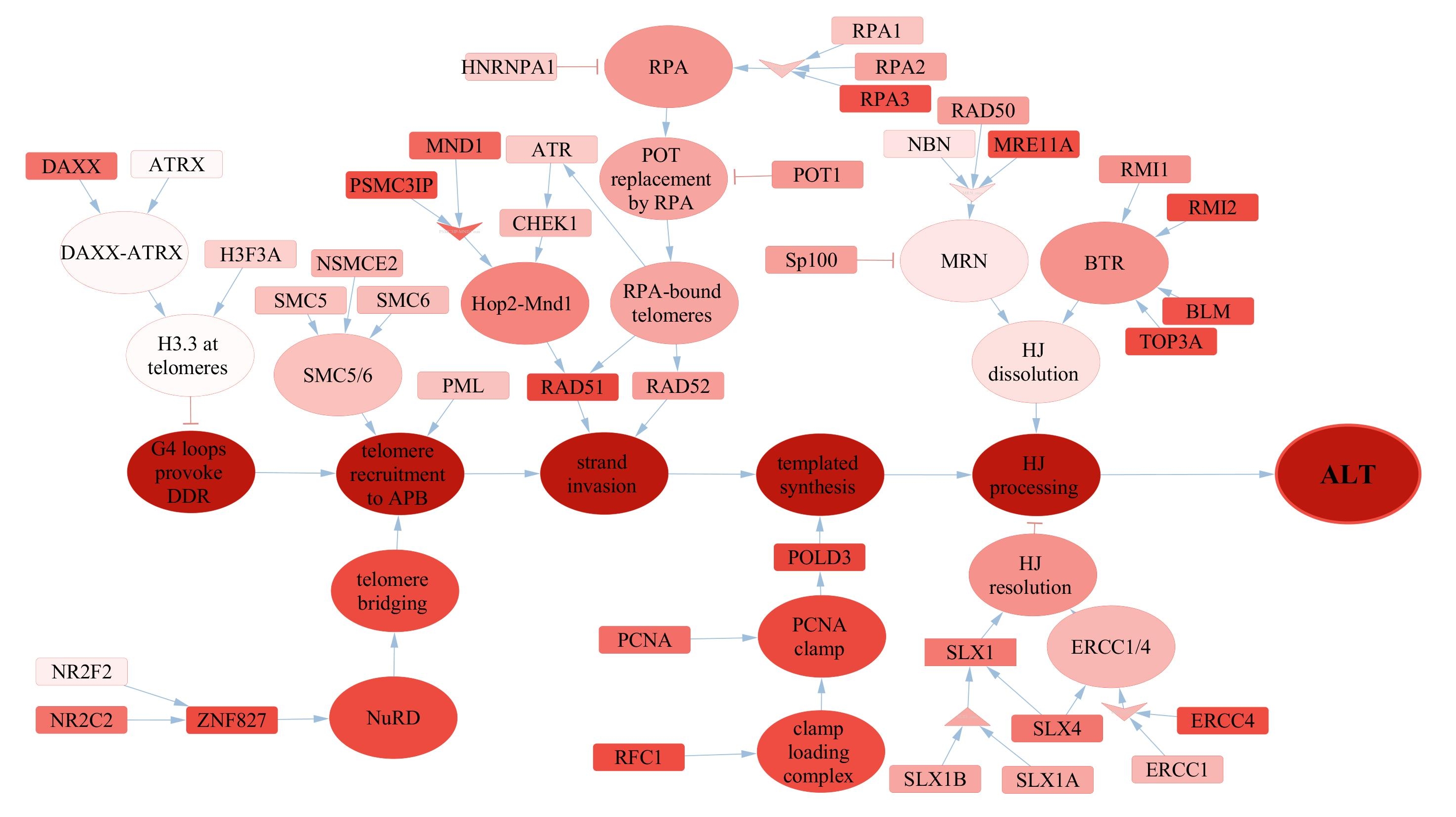

### ALT_A7.jpg

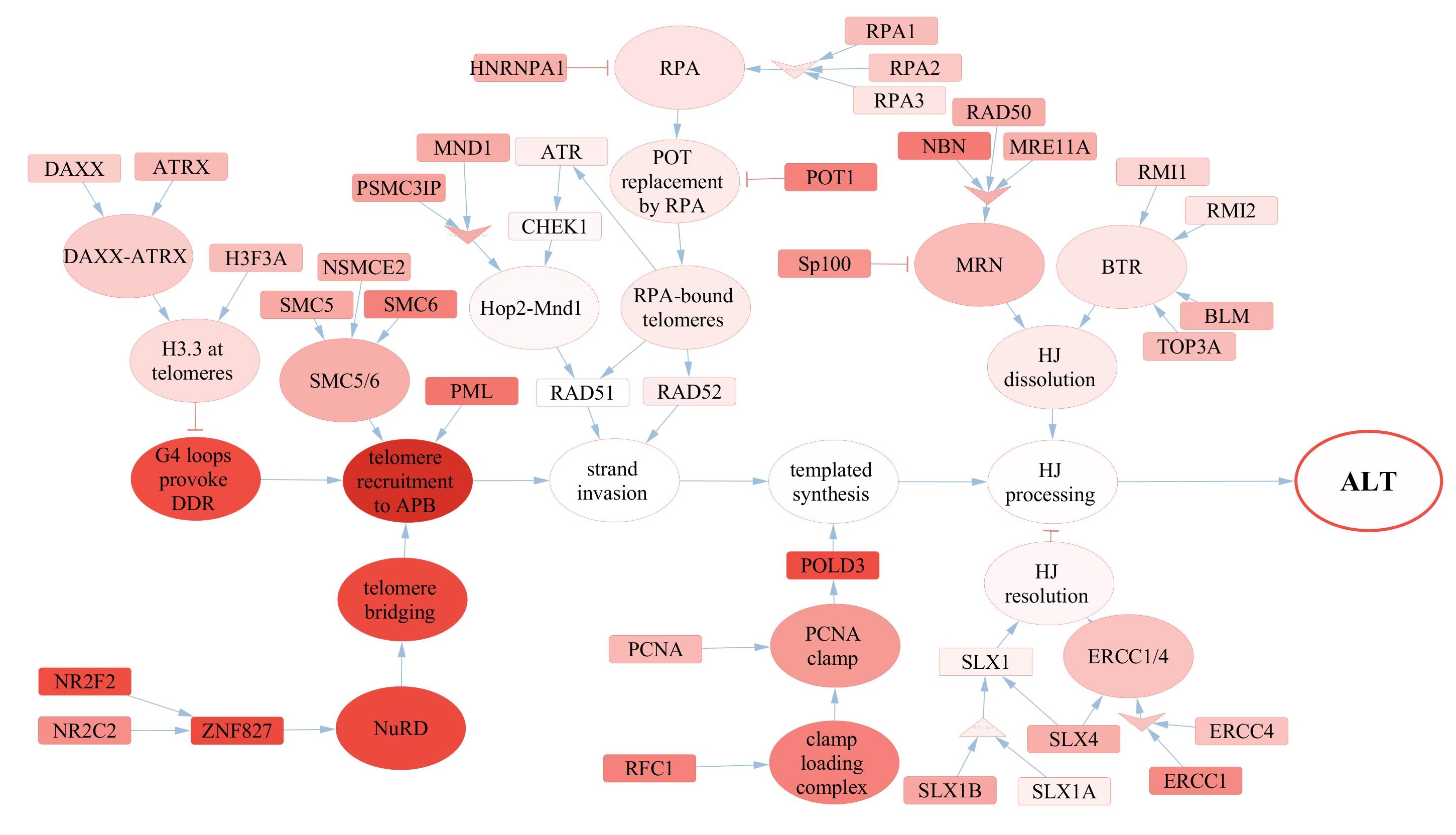

### ALT_C33_1.jpg

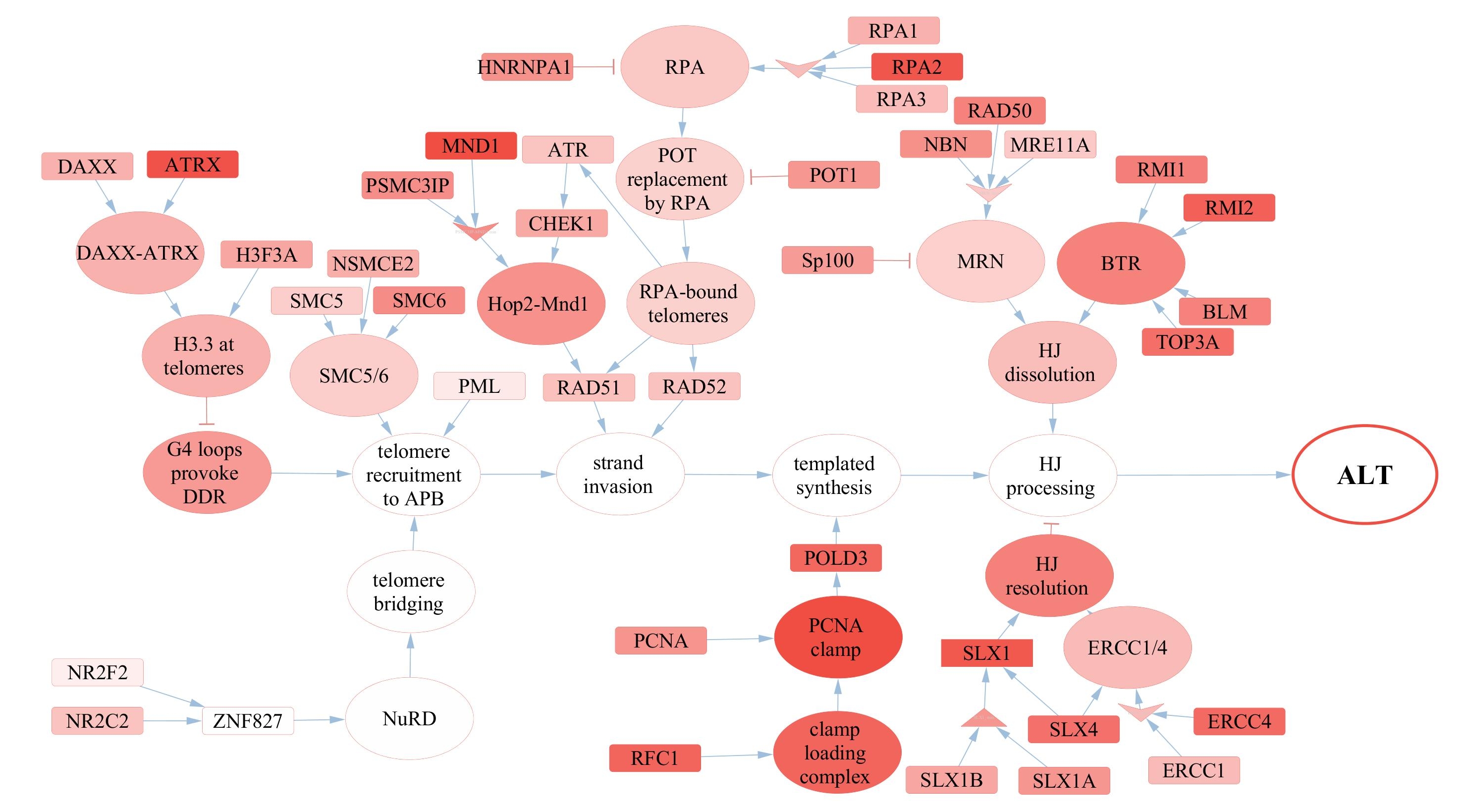

### ALT_IMR90_1.jpg

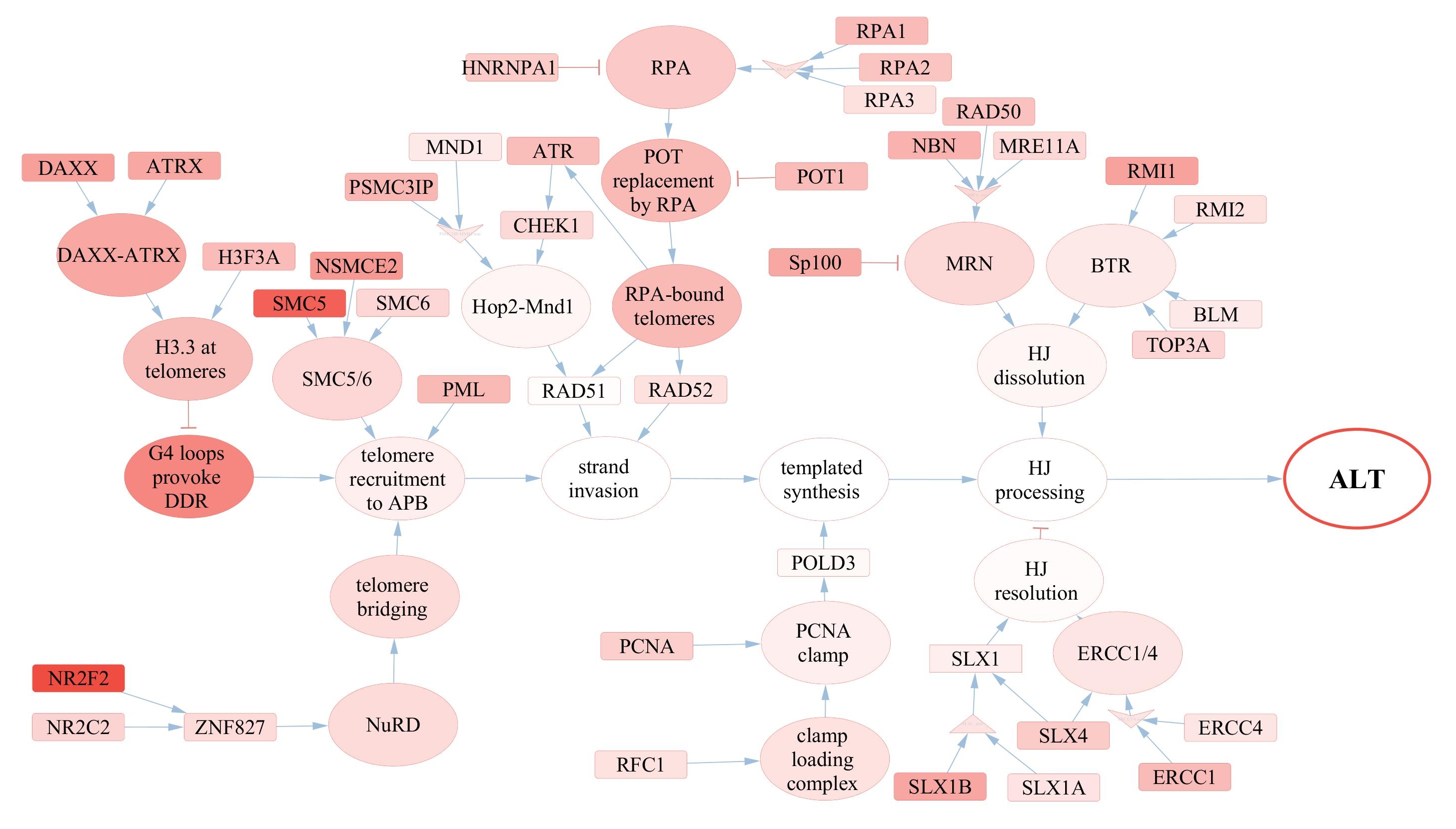

### ALT_SUSM1_1.jpg

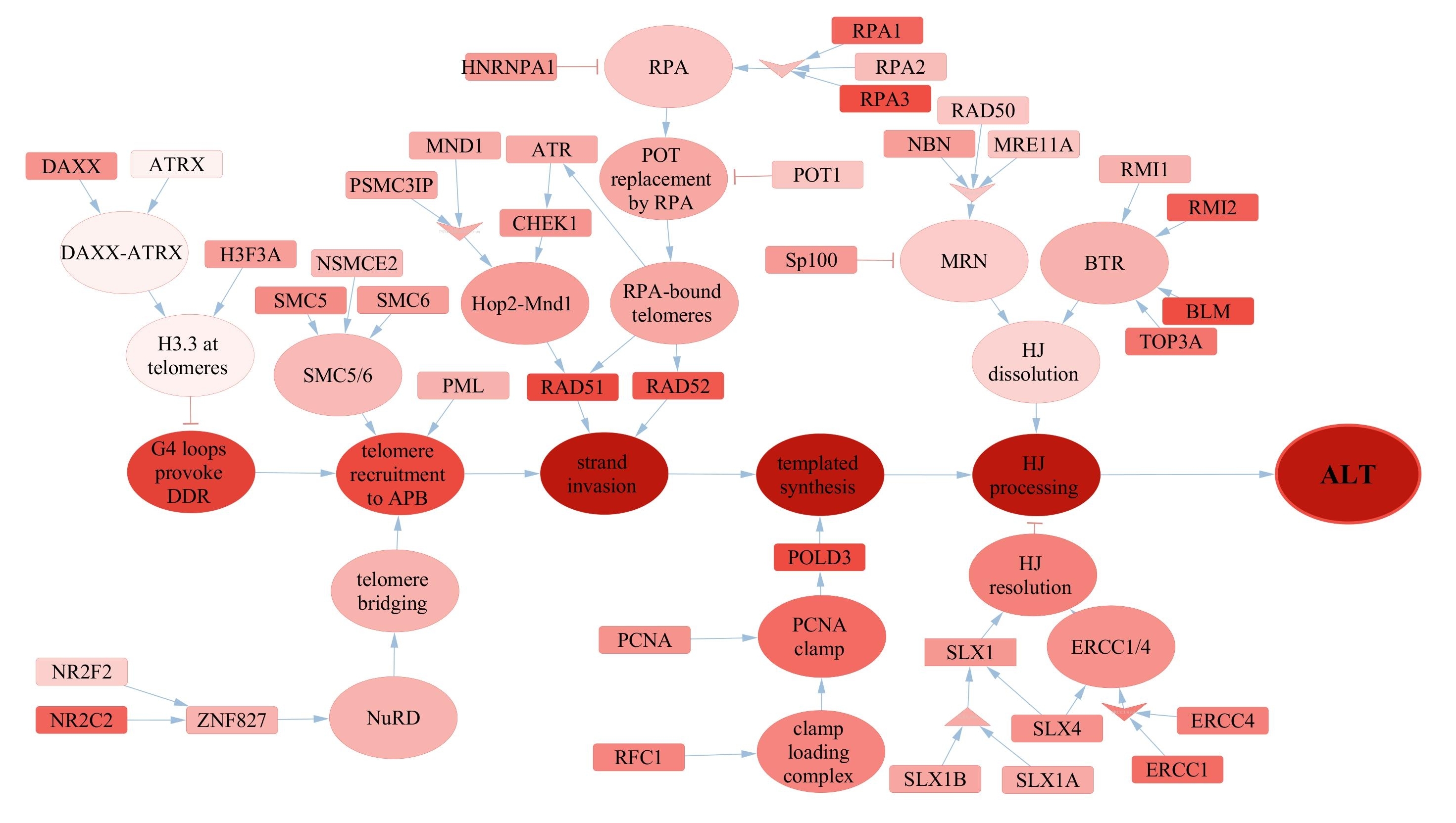

### ALT_T5.jpg

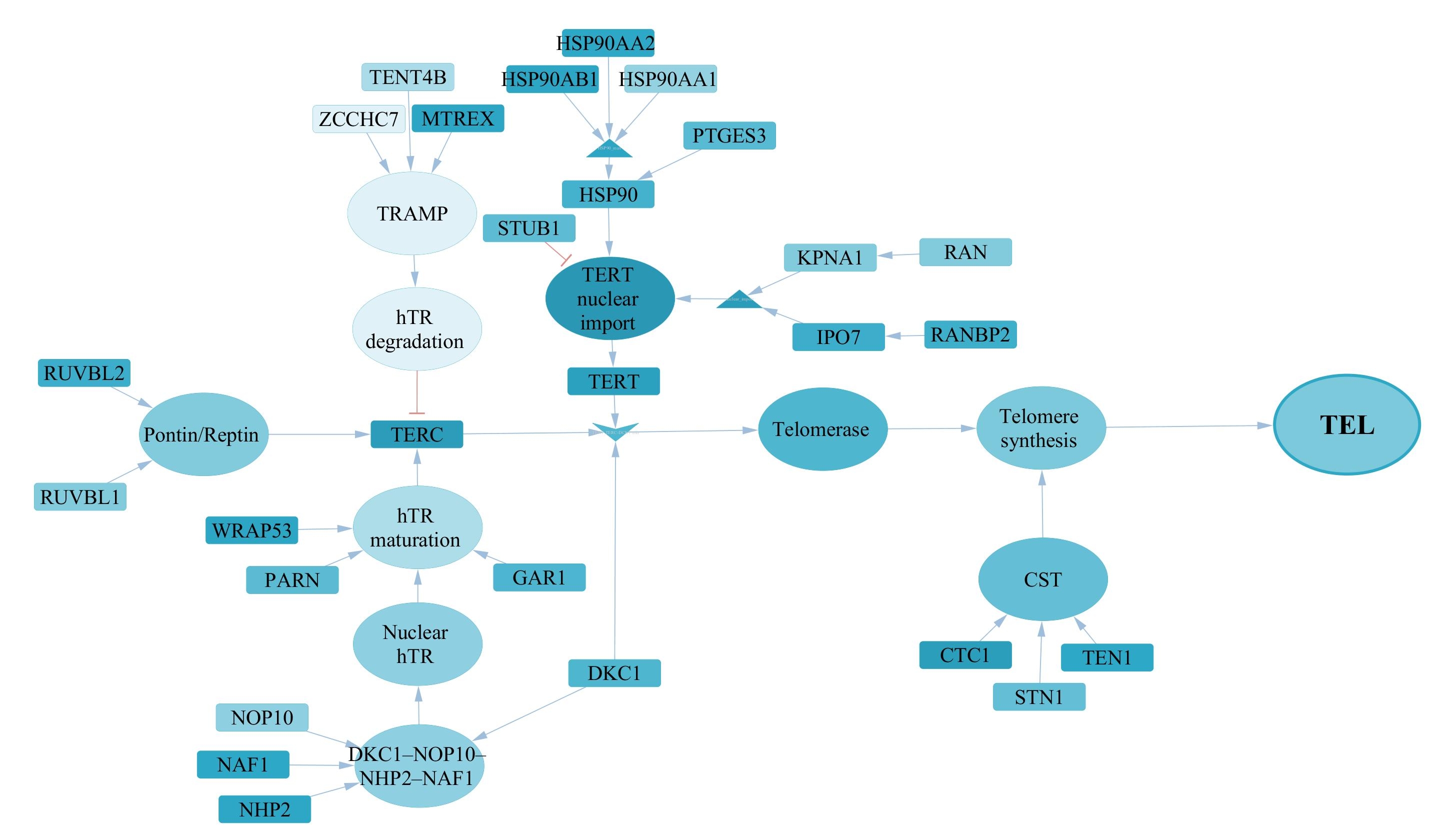

### ALT_T8.jpg

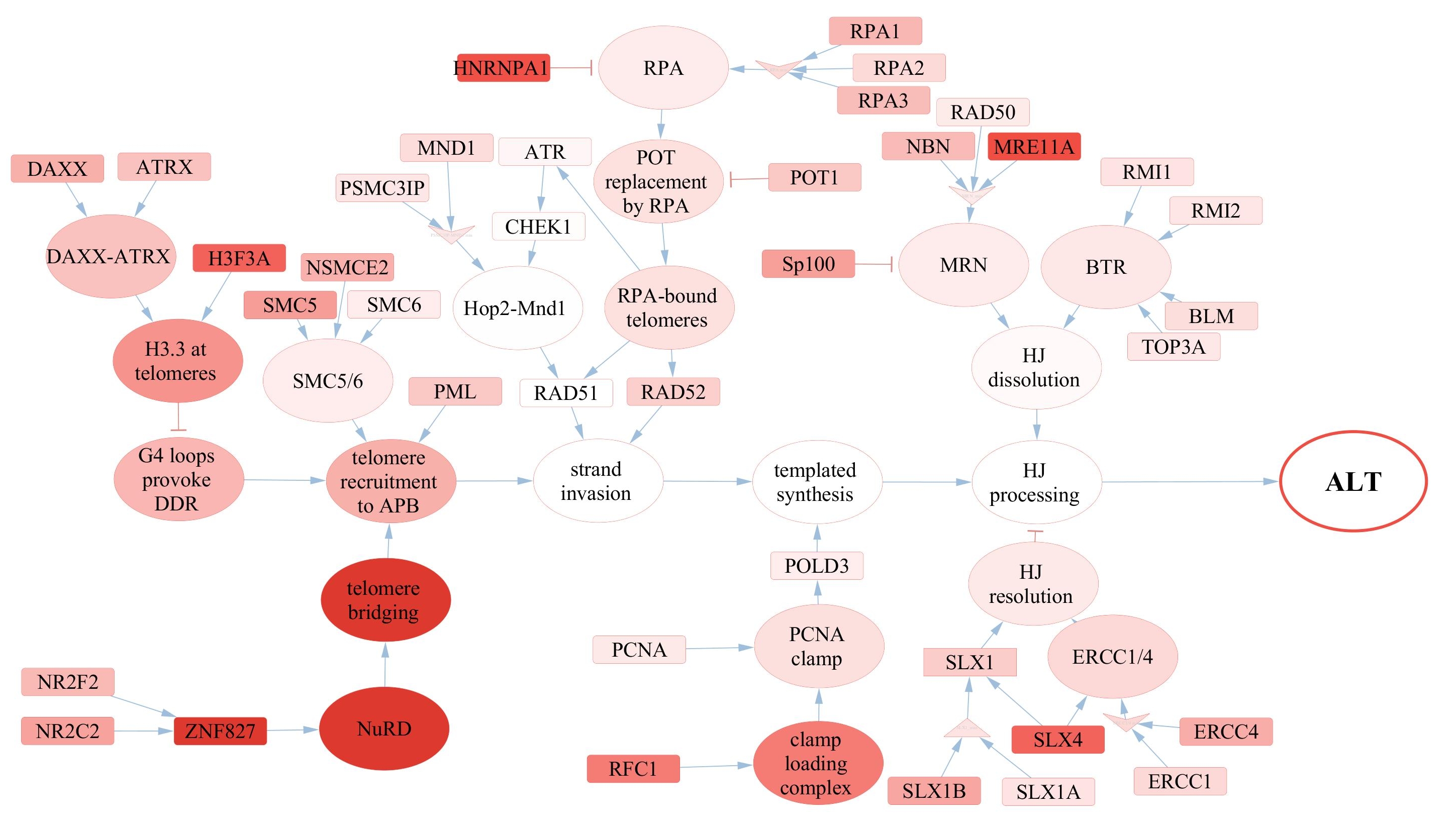

### TEL_5637_1.jpg

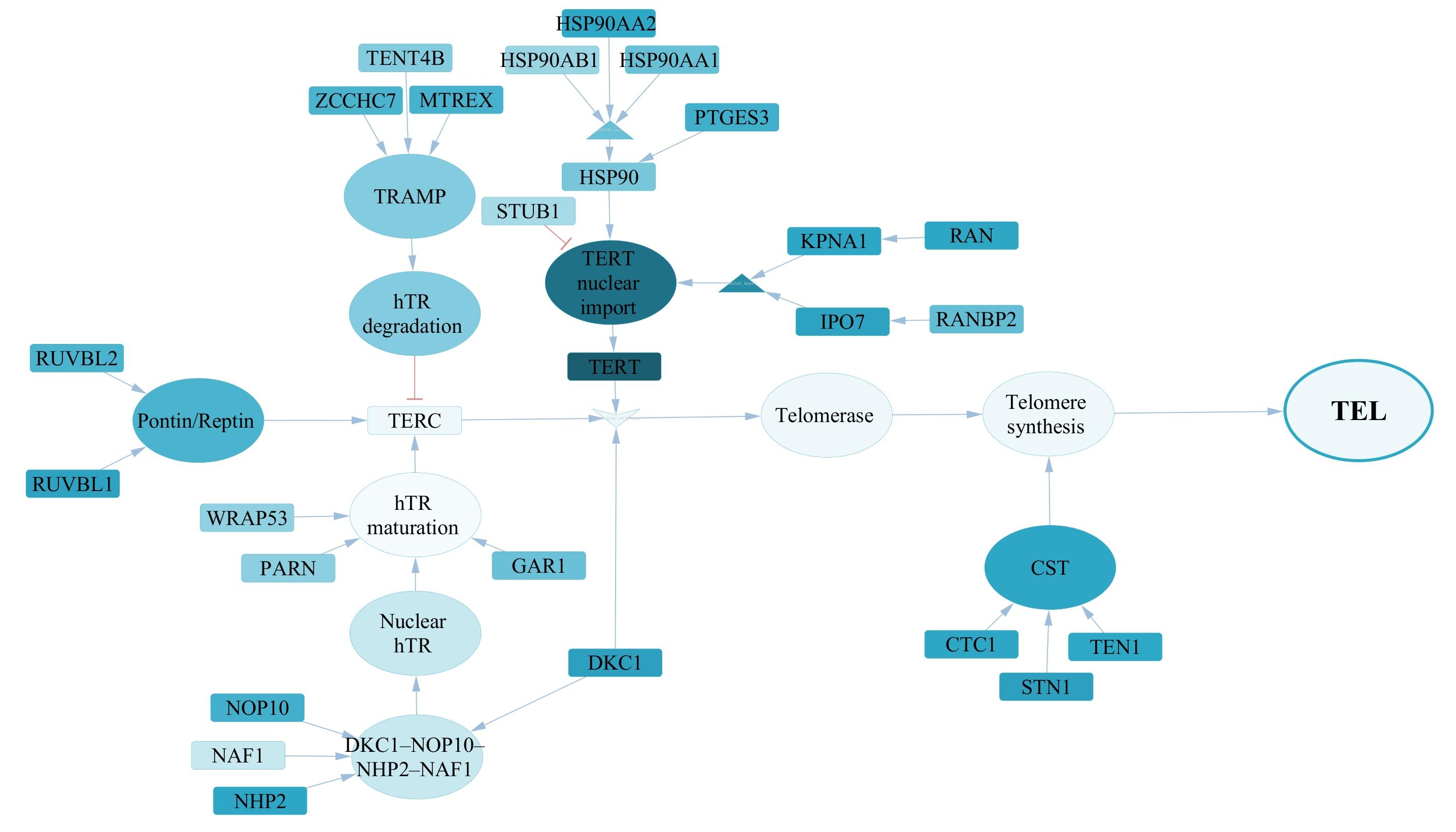

### TEL_A1.jpg

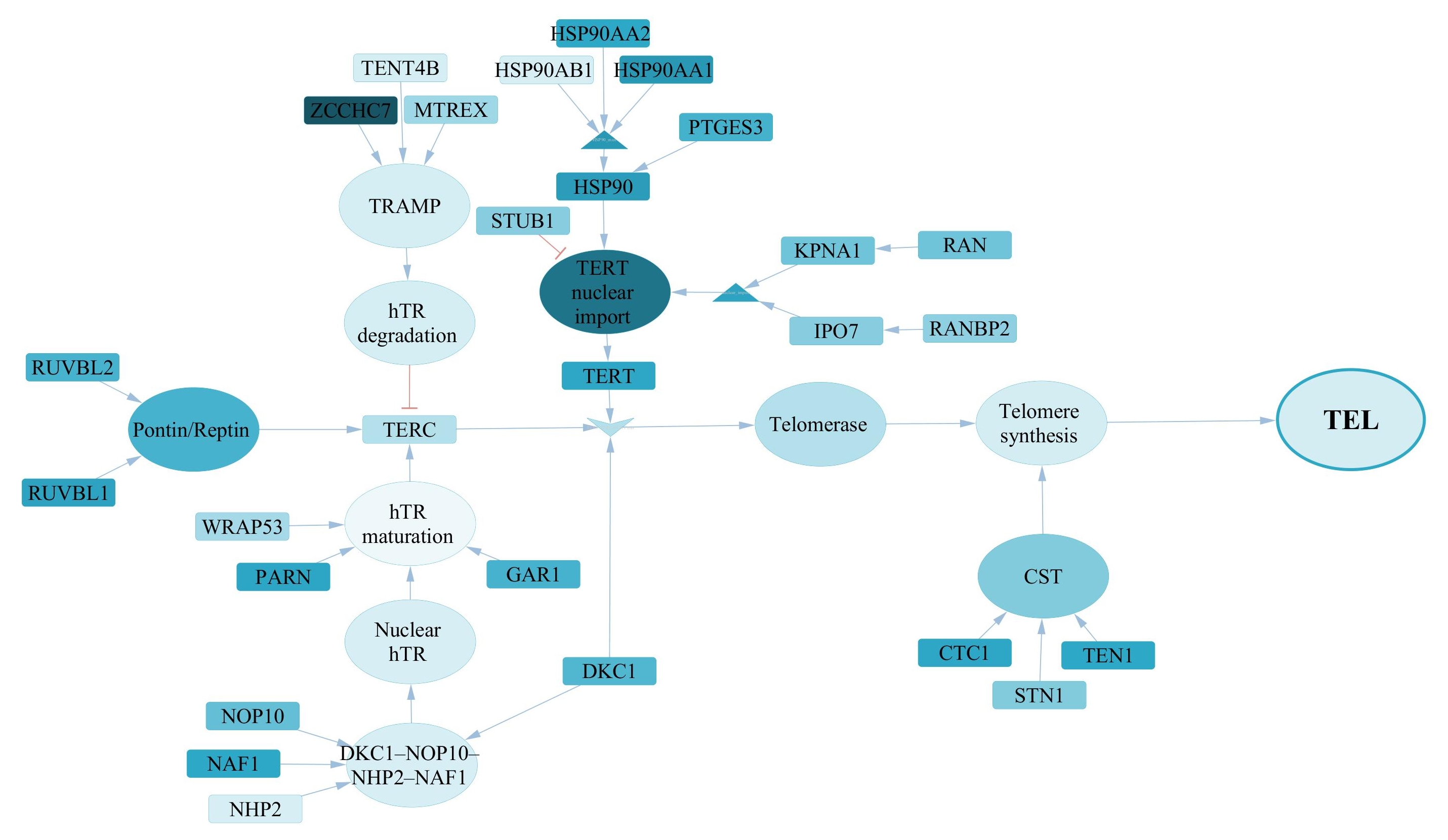

### TEL_A7.jpg

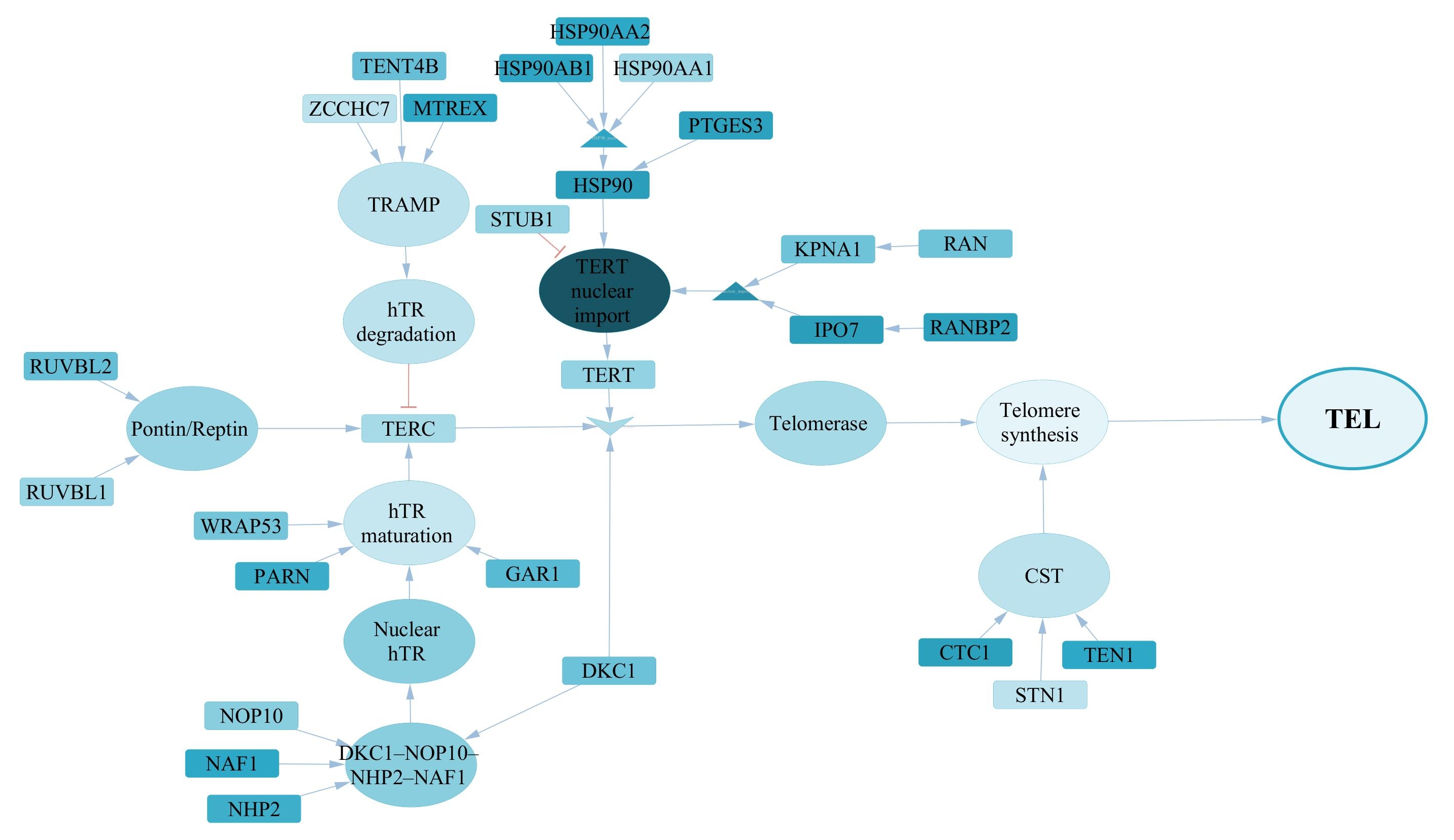

### TEL_C33_1.jpg

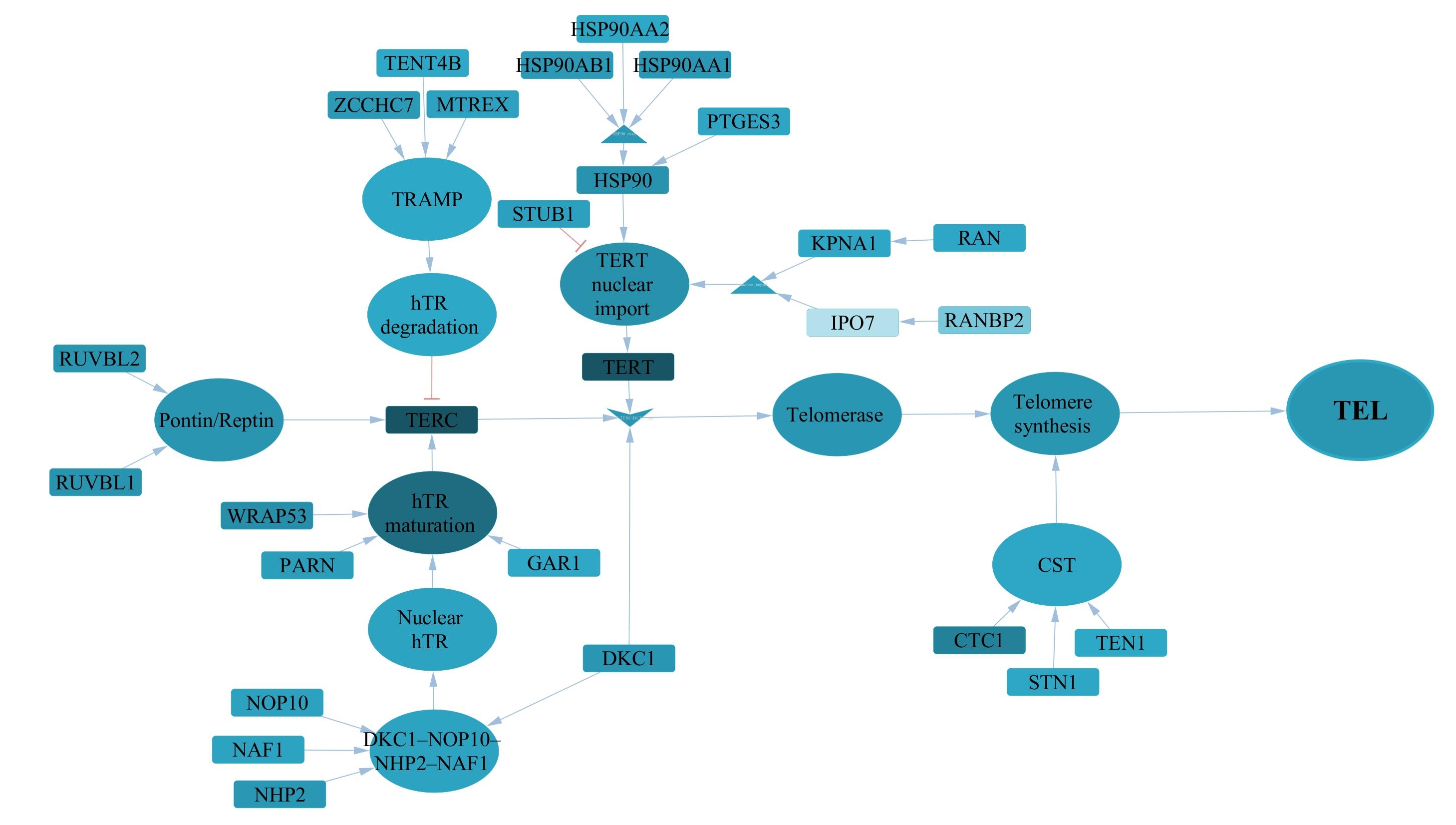

### TEL_IMR90_1.jpg

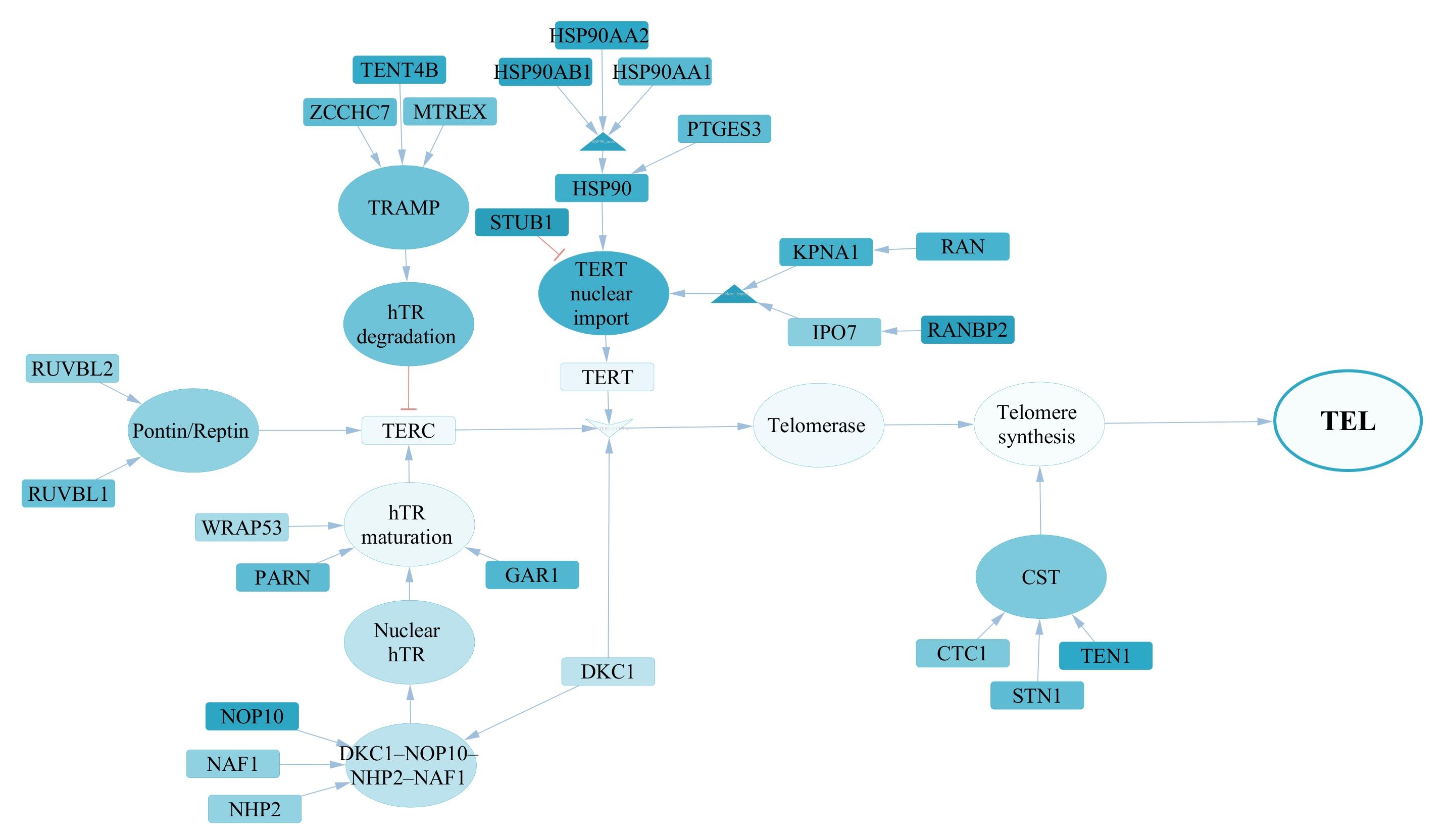

### TEL_SUSM1_1.jpg

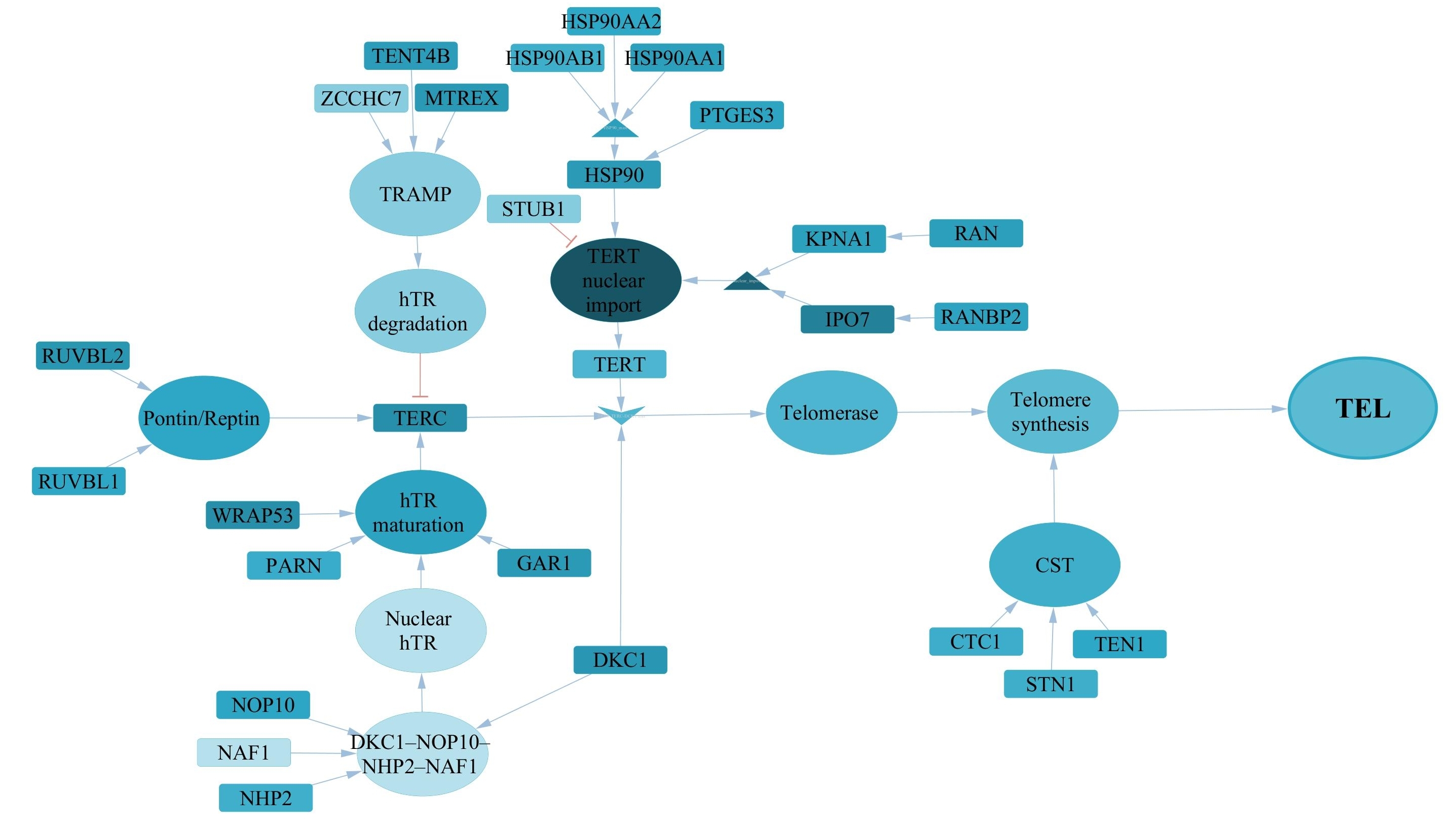

### TEL_T5.jpg

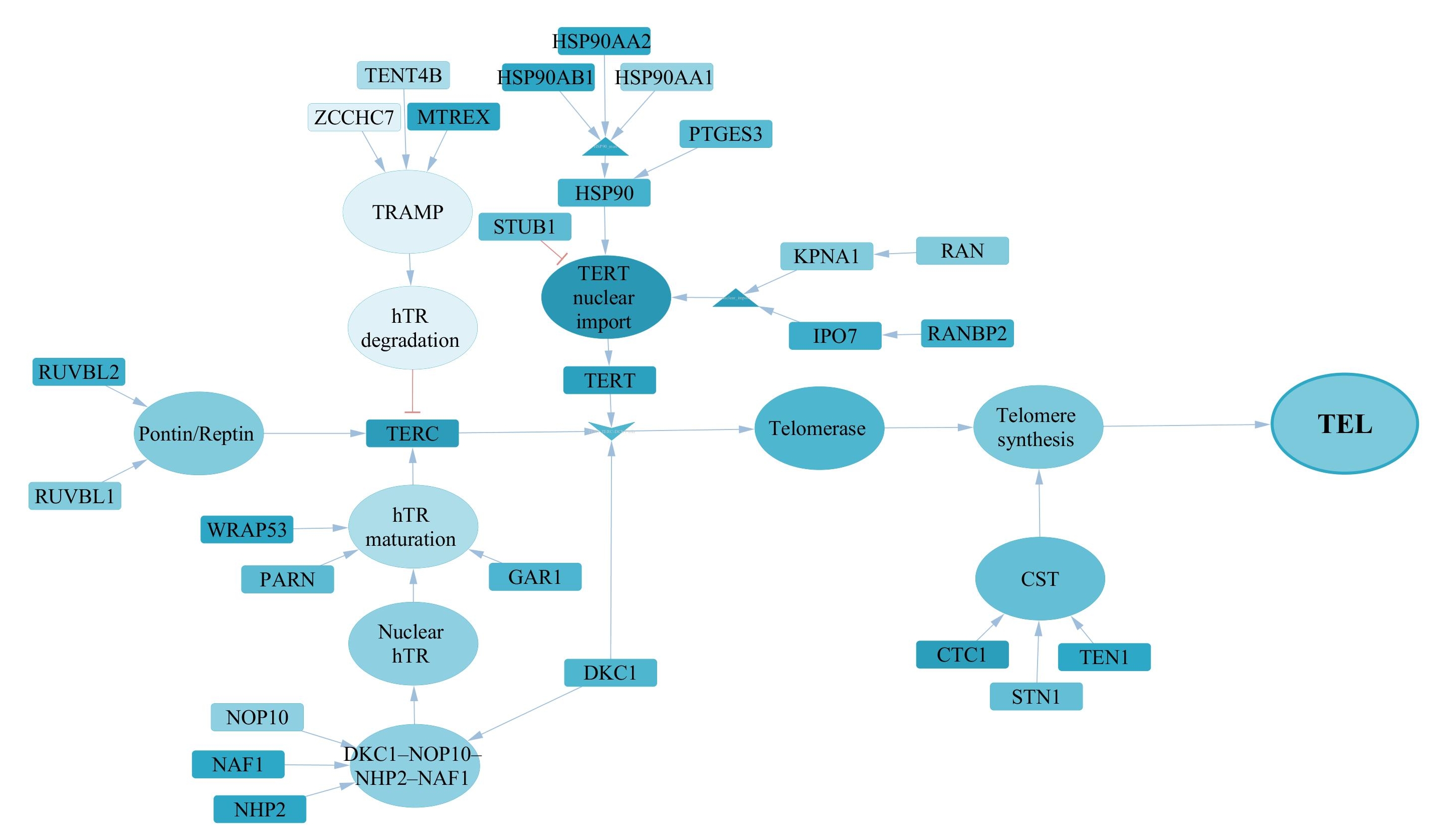

### TEL_T8.jpg

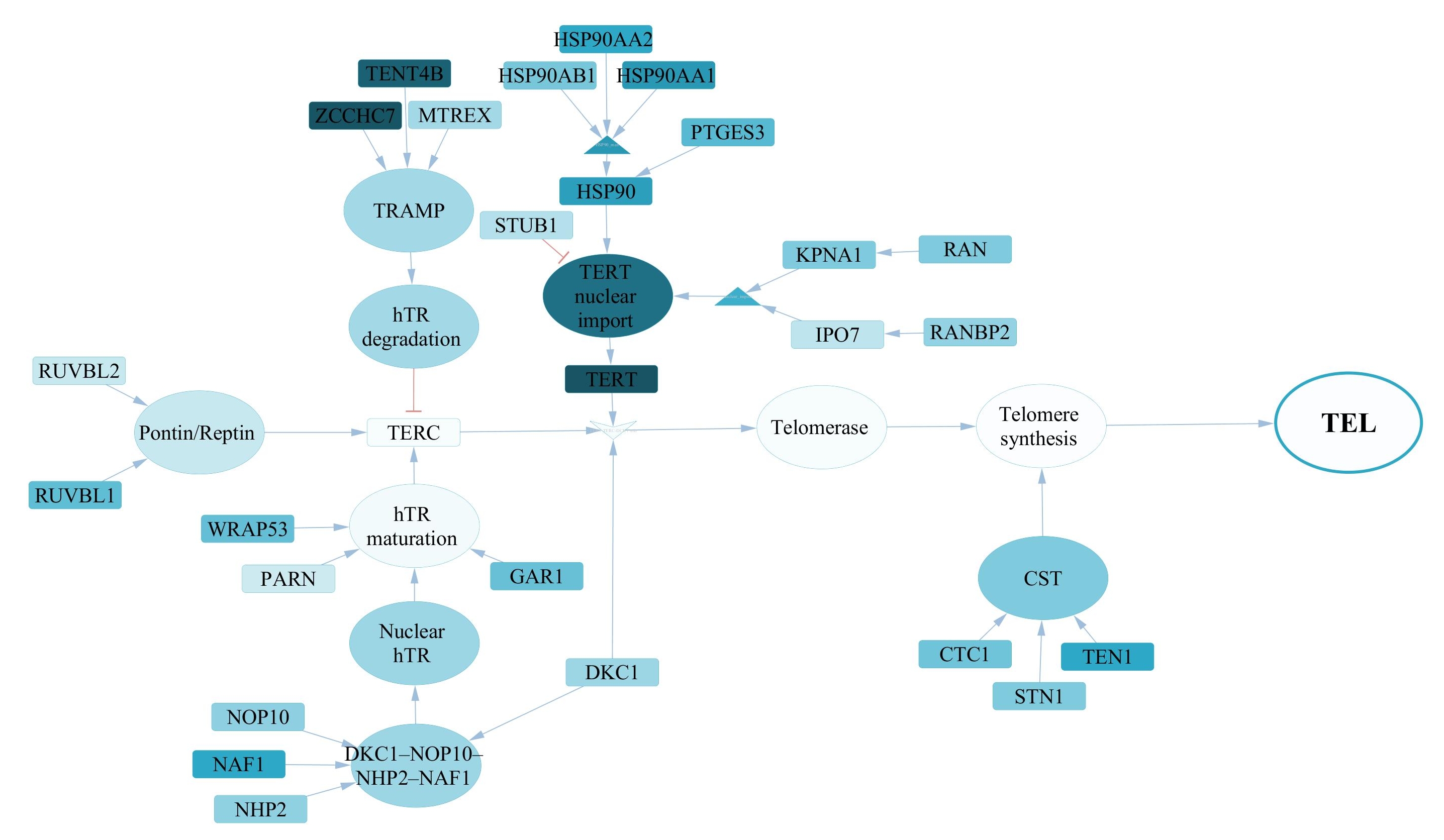
